# Coordinated regulation of photosynthesis and translation via NIK1/RPL10/LIMYB signaling module in response to biotic and abiotic stresses

**DOI:** 10.1101/2023.06.13.544461

**Authors:** Ruan M. Teixeira, Marco Aurélio Ferreira, Otávio J.B. Brustolini, Thainá F.F. Saia, James Jean-Baptiste, Samera S. Breves, Igor N. Soares, Nathalia G.A. Ribeiro, Christiane E. M. Duarte, Lucas L. Lima, Leandro Licursi Oliveira, Humberto J.O. Ramos, Pedro A.B. Reis, Elizabeth P. B. Fontes

## Abstract

Photosynthesis and translation are targets of metabolic control and development in plants, yet, how stress signals coordinately regulate these opposing energy-producing and consuming processes remains enigmatic. Here, we described a growth control circuit that ties the photosynthetic function to translational control in response to biotic and abiotic signals. We showed first that the downstream component of the NIK1/RPL10 antiviral signaling module, LIMYB, which represses translational machinery-related gene expression and translation, also suppresses photosynthetic apparatus-related genes leading to inhibition of the photosynthetic function. The repressing transcriptional activity of LIMYB, which was regulated by phosphorylation, was the primary determinant for the decrease in electron transport rate, exchange gas parameters, quantum efficiency, and water-use efficiency in the LIMYB-overexpressing lines. The decreased photosynthetic activity was linked to the NIK1 antiviral signaling and stunted growth. NIK1 activation by viral or bacterial PAMPs, or expressing a constitutively activated NIK1 mutant, T474D, repressed the photosynthesis-related marker genes and inhibited the photosynthetic function in control lines but not in *lymyb*. We also showed that heat and osmotic stress activate the NIK1/RPL10/LIMYB signaling circuit readouts in wild- type lines. Conversely, in *limyb-32* knockout, heat and osmotic stress induced NIK1 phosphorylation but did not cause repression of the marker genes, indicating that LIMYB links NIK1 activation to the stress-mediated downregulation of translation- and photosynthesis-related genes. The coordinated repression of photosynthesis and translation via the stress-activated NIK1/RPL10/LIMYB signaling module may adjust the plant growth pattern in response to the changing environment.

**Short summary:** The receptor-like kinase NIK1 (NSP-Interacting Kinase 1) undergoes phosphorylation under multiple biotic and abiotic signals activating the NIK1/RPL10/LIMYB signaling circuit, which coordinately downregulates translation and photosynthesis in response to the changing environment.

## INTRODUCTION

Plants are constantly exposed to environmental changes that can impact their growth, including changes in temperature, water supply, and attacks by pathogens and insects. However, understanding how these environmental changes specifically influence plant growth control is highly complex task. The existing knowledge about the underlying mechanisms is still fragmented and rudimentary making it challenging to fully assemble the molecular puzzle that governs plant growth under stress conditions. In plants, two crucial processes that promote cell growth, namely translation and photosynthesis, are profoundly affected by stress conditions (Reinbothe et al., 2010; Muhammad et al., 2021; Song et al., 2021a). In addition, the activation of defense responses to pathogens often provokes stunted growth (Song et al., 2021a). Despite these observations, we have only limited understanding about the molecular mechanisms that connect abiotic and biotic stress signalswith the coordinated regulation of photosynthesis and translation.

The energy for plant growth is obtained via respiration and photosynthesis. Photosynthesis not only directly provides substrates for the ATP-generating mitochondrial reactions but also supplies C skeletons for allocation in protein synthesis. Any stress condition that interferes with energy and carbon balance in the photosynthetic cell is expected to affect translation (Song et al., 2021a). Consequently, the photosynthesis rate correlates with carbon allocation to translation, polysome abundance, and translational regulatory processes, including phosphorylation of ribosomal proteins (RP) and translation initiation factors (Tcherkez et al., 2020). While the impact of photosynthesis on translation has been established at some level, the molecular mechanisms underlying the coordinate regulation of these processes are still poorly understood.

As the most energy-consuming cellular process, translation is often tightly regulated under stress conditions. A well-established mechanism for translational control in plant cells involves the TOR signaling pathway, which serves as a master regulator of cell growth (Burkart and Brandizzi, 2021). This conserved pathway, found in both mammalian and plant cells, senses various factors, including nutrient availability, energy status, hormones, and stress signals. Therefore, it governs essential processes, including ribosome biogenesis, translation, and transcription of a set of translation machinery-related genes (Reinbothe et al., 2010). As a nutrient-sensing pathway, the TOR kinase is activated in response to the availability of sugar produced by photosynthesis and light (Song et al., 2021b; Han et al., 2022). In addition to its role in translation regulation, TOR signaling also influences photomorphogenesis and plays a regulatory role in leaf and chloroplast development.

Another translational control mechanism associated with plant immunity is the NUCLEAR SHUTTLE PROTEIN (NSP)-INTERACTING KINASE 1 (NIK1)-mediated antiviral signaling (reviewed in Machado et al., 2015; 2019; Fontes et al., 2021; Teixeira et al., 2021; Ferreira et al., 2021). NIK1 is a leucine-rich repeat receptor-like kinase (LRR-RLK) iniitially identified as a virulence target of the begomoviral NSP (Fontes et al., 2004: Mariano et al., 2004). Upon viral infection or recognition of viral pathogen-associated molecular patterns (PAMPs; begomovirus-derived nucleic acids), NIK1 undergoes oligomerization with itself or a yet-to-be-identified viral pattern recognition receptor (PRR), resulting in phosphorylation at a critical threonine residue, position 474 (Santos et al., 2009; Teixeira et al., 2019). Activated NIK1 then mediates the phosphorylation of the ribosomal protein L10, redirecting RPL10 to the nucleus (Carvalho et al., 2008). In the nucleus, phosphorylated RPL10 interacts with the transcriptional repressor L10-Interacting Myb domain-containing protein, LIMYB, resulting in the full repression of translational machinery-related genes, including RP genes and translational initiation and elongation factors (Zorzatto et al., 2015). Prolonged activation of NIK1 leads to suppression of global translation (Zorzatto et al., 2015). Begomovirus cannot escape this translational regulatory mechanism of plant cells; the viral mRNAs are not efficiently associated with polysomes and translated, compromising infection (Brustolini et al., 2015; Zorzatto et al., 2015). Viral NSP binds to the kinase domain of NIK1 to counteract this antiviral defense (Fontes et al., 2004; Santos et al., 2009). Therefore, the NIK1-mediated antiviral signaling is evolutionarily overcome by the viral suppressor NSP. Nevertheless, NIK1 antiviral signaling also cross-communicates with antibacterial immunity imposing relevant implications (Li et al., 2019). Firstly, under normal conditions, NIK1 interacts with the pattern recognition receptor (PRR) FLS2 and its coreceptor BAK1 to prevent autoimmunity and hence interference with growth. Secondly, NIK1 interaction modulates the formation of the PAMP (flagellin)-induced FLS2-BAK1 immune complex, thereby controlling the extent of PAMP-triggered immunity (PTI) activation that could otherwise impact growth. Finally, the activated immune complex FLS2-BAK1 phosphorylates NIK1 at the key threonine-474 residue initiating the transduction of an antiviral signal that ultimately results in global translation suppression. Therefore, bacterial infection may induce NIK1-dependent resistance against subsequent virus infections. The bacterial PAMP-induced activation of NIK1-mediated host translational suppression may also contribute to the stunted growth observed during immune responses.

Recent network-centric analyses of the LRR-based cell surface interaction network (CSILRR) have provided evidence suggesting that NIK1 may be one of the most influential LRR-RLKs as information spreaders (Machado et al., 2015; Ahmed et al., 2018; Smakowska-Luzan et al., 2018; Li et al., 2019). Therefore, NIK1 may oligomerize with different stimulus-sensing receptors to control translation under adverse conditions. We tested this hypothesis using two different approaches. Firstly, we uncovered the transcriptional landscape resulting from NIK1 activation by performing a ChiP-seq on LIMYB-GFP seedlings. Our objective was to examine whether NIK1 activation is associated with cellular processes beyond translation, thus providing insights into potential broader functions of NIK1. Secondly, we monitored the NIK1 activation under different stimuli known to elicit shared responses. By subjecting the plant system to these stimuli, we aimed to evaluate the extent to which NIK1 activation could be modulated and synchronized across diverse signaling pathways.These experimental approaches were undertaken to deepen our understanding of NIK1’s role in signal transduction and translation regulation, and to shed light on its potential interactions with other receptors and cellular processes.

## Results

### LIMYB represses the promoter activity and expression of photosynthetic apparatus-related genes leading to photosynthesis inhibition

As a downstream component of the NIK1 antiviral signaling, LIMYB represses translational machinery-related genes, which accounts for the NIK1-mediated translational control mechanism (Zorzatto et al., 2015). To assess further the LIMYB regulon, we performed an RNA-sequencing (RNA-seq) and chromatin immunoprecipitation sequencing (ChIP-seq) experiments on LIMYB-GFP-overexpressing seedlings (Supplemental Figure 1A and 1B). Through the integration of RNA-seq and ChIP-seq data, we identified a set of differentially expressed genes that were directly regulated by LIMYB. RNA-seq was performed with Col-0, LIMYB32-L1, and LIMYB32-L3 RNA (Supplemental Figure 1C and 1D). The LIMYB-induced transcriptome was represented by 1105 upregulated genes and 1246 downregulated genes resulting in a total of 2351 differentially expressed (DE) genes (>2-fold change up or down, *P* < 0.01, Supplemental Figure 1E and 1F).

The top (ranked by false discovery rate [≤0.05]) significantly enriched GO term was structural constituents of ribosomes, supporting the role of LIMYB in repressing translation-related processes (Supplemental Figure 1E; Supplemental Figure 2; Zorzatto et al., 2015). The ChIP-seq analysis identified LIMYB binding peaks across the Arabidopsis genome, resulting in 6248 represented transcripts. By overlapping these transcripts with the RNA-seq results, we identified 204 upregulated and LIMYB-bound genes, as well as 156 downregulated and LIMYB-bound genes (Supplemental Figure 3A and 3B). These overlapping genes constituted the core LIMYB up-regulon and down-regulon, respectively. The enriched down- and upregulated categories of genes within these regulons (Supplemental Figure 3A and 3B) are described in Supplemental Tables 1 and 2. Notably, these regulons included several previously known LIMYB-directly regulated genes, such as ribosomal protein (RP) genes (*RPL18E, RPL28E, RPS13*A, *RPS25*, *RPL13*, *RPL4/L1),* as well as newly identified LIMYB targets (Supplemental Table 3; Zorzatto et al., 2015).

Consistent with its role in translational control, the core LIMYB down-regulon exhibited significant enrichment of translation-related GO terms, including structural constituent of ribosomes and translation (Supplemental Figure 3A; Supplemental Table 1). The RNA-seq and ChIP-seq results indicated that several translational regulatory genes (translational initiation and elongation factors) and RP genes were LIMYB-regulated targets (Supplemental Tables 3 and 4). *De novo* discovery of enriched motifs within the LIMYB binding peaks identified the MYB-related binding sites C[A/C/T]AA[A/C/G]C (E-value 3.0e-350) and AAGAA[A/G][A/C] (E-value 3.8e-252) and a conserved element [A/G]TATAT[A/G] (E-value 1.0e-0.97) as top-scoring motifs (Figure 1A). To examine functional significance of this motifs as cis-regulatory elements for LIMYB assembly on promoters, we mapped the ChIP-seq peaks and RNA-seq hits to the loci of selected RP genes, and translational regulatory factors (Figure 1B). The ChIP-seq peaks were mapped to these motifs on the promoter region of these LIMYB target genes. The corresponding transcripts were downregulated in the LIMYB-overexpressing lines, as shown by the RNA-seq hits and confirmed by RT-qPCR (Figure 1B and Supplemental Figure 2B). Additionally, a promoter transactivation assay demonstrated that a truncated version of the RPL18A promoter (pL18A, 325 bp) lacking the LIMYB binding sites at −455 position (Supplemental Figure 3D) was no longer downregulated by LIMYB in Arabidopsis protoplasts (Figure 1C). The accumulation of LIMYB transgene in the protoplast was monitored by immunoblotting (Supplemental Figure 3C).

**Figure 1.**
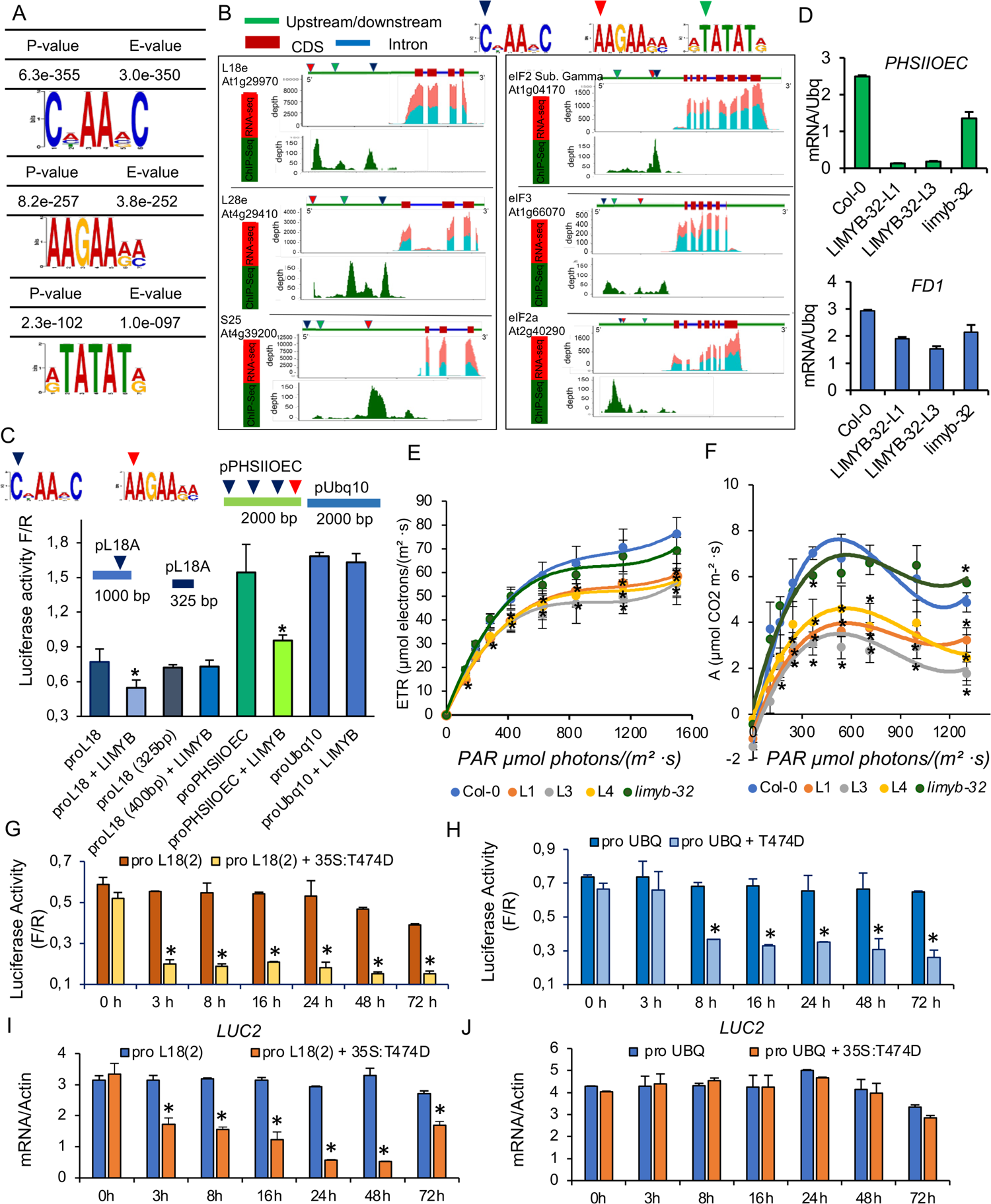
LIMYB represses the expression of photosynthetic apparatus-related genes and inhibits photosynthesis. **(A)** Top enriched motifs in the promoter region of LIMYB-regulated genes. **(B)** LIMYB DNA-binding motifs are enriched in the promoter region of LIMYB downregulated genes. In the schematic representation of the loci indicated in the figure, upstream promoter sequences are green, and coding regions are red. ChIP-seq peaks map to the promoter region of the indicated LIMYB-repressed genes, whereas the RNA-seq hits lay in the coding region. The relative expression of the indicated gene in Col-0 and LIMYB-overexpressing lines is shown by the abundance of RNA-sequencing hits in blue (LIMYB) and red (Col-0). **(C)** LIMYB represses the PSIIOEC promoter. *N. benthamiana* leaves were agroinfiltrated with plasmids carrying the promoter length (as indicated in the figure) fused to luciferase and the 35S: LIMYB construct. After 48h, luciferase activity was measured from total protein extracts of transformed leaves. An unrelated Ubiquitin promoter (UBQ) was used as a negative control. Results are mean values ±SD (n=6). Asterisks indicate significant differences from the control line (t-test, p<0.05). **(D)** *PHSII-OEC* (Photosystem II oxygen-evolving complex) and *FD1* (Ferredoxin) genes are repressed by LIMYB. The transcript levels of the indicated genes were quantified by RT-qPCR in *LIMYB*-overexpressing (LIMYB-32-1 and LIMYB-32-L3) lines and the wild-type (Col-0) line. Asterisks indicate significant differences from the control line (t-test, p<0.05). **(E)** Electron transport rate (ETR) in *LIMYB*-overexpressing lines (LIMYB-32-L1, L3, L4), and *limyb* knockout line. PAR denotes Photosynthetically Active Radiation. **(F)** CO_2_ assimilation rate (A) is inhibited in *LIMYB*-overexpressing lines. **(G)** The constitutively activated NIK1 mutant T474 represses the RPL18 promoter activity. Arabidopsis protoplasts were transformed with the proL18:luciferase construct alone or combined with the 35S:T474 construct. Luciferase activity was determined at different time points. **(H)** T474 does not affect the activity of the UBQ-unrelated promoter. Arabidopsis protoplasts were transformed with the proUBQ:luciferase construct alone or with the 35S:T474 construct. Luciferase activity was determined at different time points. **(I)** T474D represses the RPL10 promoter activity. Arabidopsis protoplasts were transformed with the proL18:luciferase construct alone or combined with the 35S:T474 construct. Transcript levels were determined by RT-qPCR. **(J)** T474D does not repress the Ubq10 promoter activity. Arabidopsis protoplasts were transformed with the proUBQ:luciferase construct alone or combined with the 35S:T474 construct, and transcript levels were determined by RT-qPCR. For g, h, i, j, the results are mean values ±SD (n=6). Asterisks indicate significant differences from the control line (t-test, p<0.05). The experiments in **C-J** were repeated at least twice with similar results.

These findings, along with previous observation of LIMYB repressing transcriptional activity and binding to RP promoters, provide further evidence for the role of LIMYB as a transcriptional repressor (Zorzatto et al., 2015). These complementary experiments implicated the core LIMYB down-regulon as direct target genes of LYMYB; thereby, shifting the focus to the core LIMYB down-regulon for more detailed analysis.

Another significantly enriched GO term in the downregulated core was represented by photosynthesis-related genes, including genes involved in photosystem II assembly, light reaction, and photosynthetic electron transport (Supplemental Figure 4; Supplemental Table 5). The enrichment of photosynthesis-related genes in the LIMYB down regulon raised the hypothesis that LIMYB coordenastes the repression of both translation and photosynthesis to balance carbon demand and allocation in response to biotic signals. Consistent with this hypothesis, responses to starvation-related GO terms were significantly enriched in the upregulated regulon (Supplemental Table 2). This hypothesis was further supported by subsequent experiments.

The expression. of selected photosynthesis-related genes, such as the *PHOTOSYSTEM II OXYGEN-EVOLVING COMPLEX* (*PHSIIOEC*) gene and *FERREDOXIN* (*FD1*) from the electron transport chain (ETC) in photosystem I, was confirmed to be repressed by *LIMYB* overexpression through RT-qPCR (Figure 1D). Furthremore, a firefly *LUCIFERASE* (fLUC) reporter construct containing a 2-kb 5’flanking region of the *PHSIIOEC* gene, which includes LIMYB binding sites, was used for transactivation assays with LIMYB in Arabidopsis protoplast (Figure 1C). The transient expression of LIMYB inhibited the PHSIIOEC promotor activity but not the UBQ promoter activity, which served as a control (Figure 1C). Finally, we showed that *LIMYB* overexpression inhibited the electron transport rate (ETR), which was associated with a decreased photosynthetic rate (*A*), stomatal conductance (*gs*), transpiration rate (*E*), internal CO2 concentration (ci), the quantum yield of PSII electron transport (ΦPSII) and water use efficiency (WUE) compared to wild-type, Col-0 (Figure 1E and 1F, Supplemental Figure 5A-E). The simultaneous decrease in ETR, gas exchange parameters (A, gs, E and Ci), ΦPSII, and WUE in LIMYB-overexpressing lines suggests that the reduced photosynthetic activity is primarily due to a lower abundance of Photosystem II, I, and ETC components rather than defects in C fixation or damages to the photosynthetic apparatus. Adjustments in the photosystem stoichiometry have been shown to affect the quantum efficiency of photosynthesis (Chow et al., 1990).

Overall, these findings support the notion that LIMYB plays a role in coordinating the repression of translation and photosynthesis, potentially to balance carbon demand and allocation under biotic signals. The regulation of photosynthesis-related genes by LIMYB was demonstrated through various experimental approaches, including gene expression analysis, transactivation assays, and measurement of photosynthetic parameters.

The LIMYB-mediated reduced photosynthetic activity was associated with stunted growth under short or long-day light regimes (Supplemental Figure 6A). Germination and root length were also lower in the LIMYB-overexpressing lines (LYMIB-32-L1 and LIMYB-32-L3) compared to the wild-type (Supplemental Figure 6B and 6C). During the reproductive stage, however, no significant differences were observed among the genotypes (Supplemental Figure 6D and 6E). Conversely, the photosynthetic function in *limyb* knockout lines did not significantly differ from the wild-type under normal conditions (Figure 1E, 1F, Supplemental Figure 5), consistent with their similar vegetative growth performance as shown here (Supplemental Figure 6) and previously (Chong et al., 2019).

We next examined whether the decreased photosynthesis in LIMYB-overexpressing lines was primarily associated with the LIMYB repressing transcriptional activity or LIMYB-mediated suppression of global translation, which would have affected the expression of the photosynthetic genes. We have previously shown that the NIK1-mediated translational suppression is a delayed response, with a 30% reduction of translation observed only after 8-h of NIK1 activation in Arabidopsis seedlings (Zorzatto et al., 2015). Here, we showed that LIMYB represses the promoter activity of *the RPL18* marker gene (Figure 1C) and induces a coordinated decrease in mRNA and protein (luciferase activity) levels after 3-h of activated NIK1-T474D expression or viral PAMP-mediated NIK1 activation (Figure 1G and 1I; Supplemental Figure 7A and 7B). In contrast, LIMYB does not affect either the UBQ promoter activity or luciferase mRNA accumulation, which remains at normal levels throughout the experimente. However, it does promote a decrease in luciferase activity 8-h after NIK1 activation, indicating the late response of translational control (Figure 1H and 1J; Supplemental Figure 7C and 7D). Notably, mRNA and activity of luciferase under the control of the LIMYB target promoter RPL18 were simultaneously reduced as an early transcriptional response to NIK1 activation, and luciferase activity did not decline further during the translational suppression phase. These results indicate that the LIMYB’s repressive transcriptional activity is the primary determinant for LIMYB-mediated control of photosynthesis.

In summary, these findings indicate that LIMYB’s role in controlling photosynthesis is primarily mediated through its transcriptional repression activity rather than its global translational suppression. The early transcriptional response of LIMYB target genes, such as RPL18, to NIK1 activation indicates that LIMYB regulates photosynthesis by directly repressing the expression of photosynthesis-related genes.

### LIMYB-mediated repression of the photosynthetic activity is coupled to NIK1 activation

We also investigated teh potential link between LIMYB-mediated control of photosynthesis and NIK1 activation. We first examined whether activation of NIK1 by viral PAMPs would lead to the repression of photosynthetic apparatus-related genes, similar to its effect on translation-related genes (Teixeira et al., 2019). Treatment of NIK1-HA seedlings with RNA (InRNA) and DNA (InDNA) from begomovirus-infected, but not from mock-inoculated (UnRNA, UnDNA), Arabidopsis Col-0 lines induced rapid NIK1 phosphorylation (Figure 2A) and resulted in *PHSIIOEC* repression (Figure 2B). This repression of *PHSIIOEC* was less pronounced in the *nik1* and *nik2* single mutants and completely abolished in the *nik1nik2* double mutant. The expression. Levels of *PHSIIOEC* and *FD1* were significantly ower in transgenic lines expressing a constitutively activated NIK1 mutant (NIK1-T474D) compared to Col-0, indicating that NIK1 signaling controls the expression of the photosynthesis-related genes (Figure 2C).

**Figure 2.**
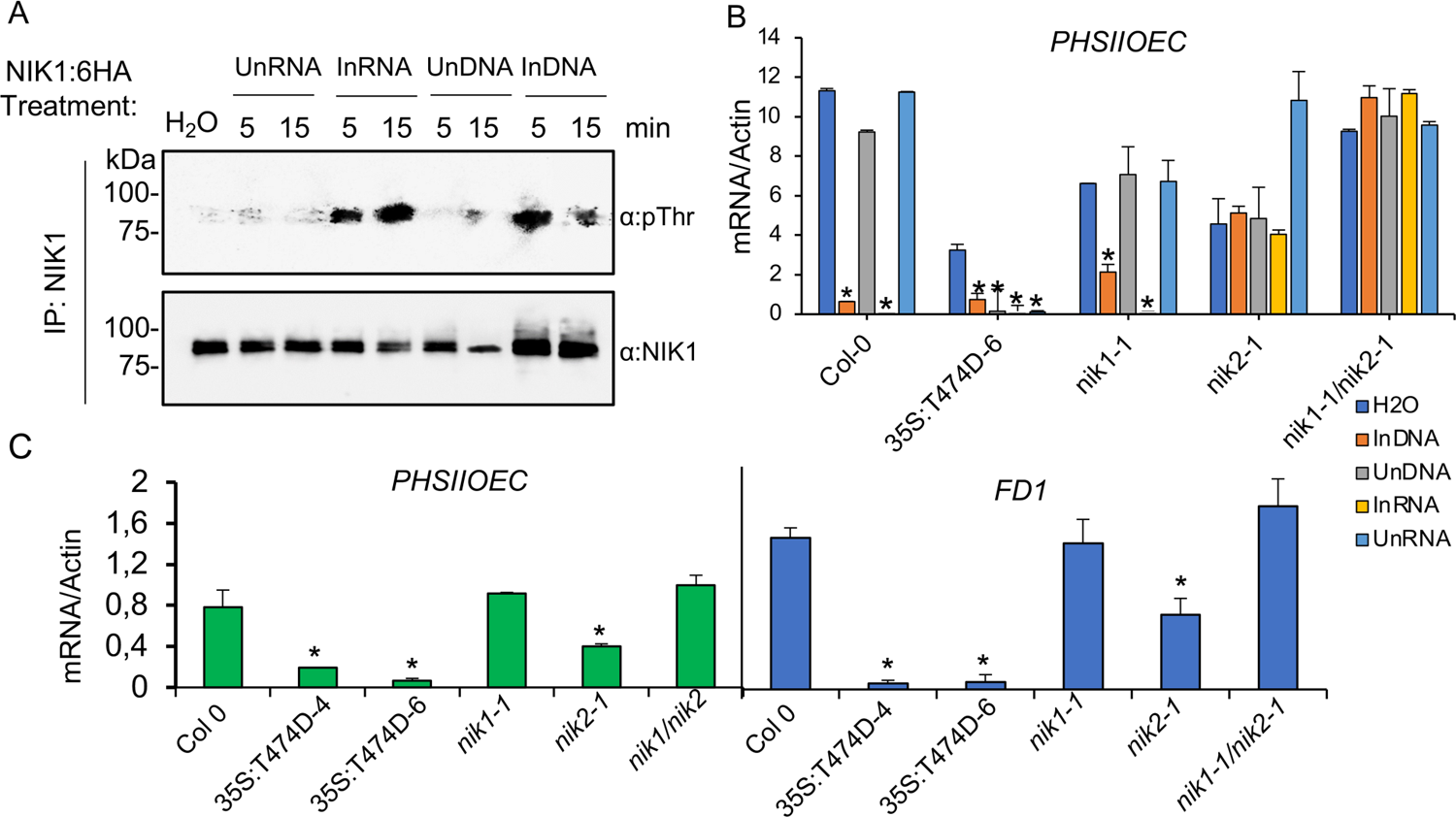
NIK1 activation by biotic signals represses the expression of photosynthetic apparatus-related genes. **(A)** NIK1 is rapidly phosphorylated by begomovirus-derived nuclei acids. NIK1-HA-expressing Arabidopsis seedlings were treated with RNA or DNA prepared from mock-inoculated (UnRNA and UnDNA) or begomovirus-infected (InRNA and InDNA) plants for the time indicated in the figure. NIK1 was immunoprecipitated with an α-NIK1 antibody and probed with an α-phospho-threonine antibody. **(B)** Viral PAMPs require NIK1 and/or NIK2 to repress the expression of the PHSII-OEC gene. Leaf discs of the indicated genotypes were treated with InRNA and InDNA, and PSII-OEC transcript levels were quantified by RT-qPCR. 35S:T474D-6 is a transgenic line that expresses the NIK1-T474D mutant. The bars represent a 95% confidence interval based on three replicates from independent experiments. Asterisks denote statistically significant differences from the control line. **(C)** Constitutive activation of NIK1 in T474D-expressing (T474D-4 and T474D-6) lines downregulates *PSII-OEC* and *Fd1*. Total RNA from the indicated genotypes and the transcript levels of the indicated genes were quantified by RT-qPCR. The bars represent a 95% confidence interval based on three replicates from independent experiments. Asterisks denote statistically significant differences from the control line. The experiments were repeated at least three times with similar results

Consistent with the NIK1 activation-mediated repression of photosynthesis-related genes, independently transformed lines expressing NIK1-T474D displayed reduced ETR compared to the control line (Figure 3), a phenotype associated with decreased gas exchange parameters, such as A, gs, E, Ci, ΦPSII, and WUE (Figure 3B-3G). Accordingly, T474 lines displayed a drastic, negative effect on growth (Supplemental Figure 8A), and exhibited lower germination and root length (Supplemental Figure 8). In contrast, the loss-of-NIK1/NIK2 function enhanced ETR, and increased the gas exchange parameters. In addition, T474D expression in *limyb* mutant did not decrease ETR and photosynthetic activity or repressed the expression of photosynthesis-related marker genes (Figure 3A-3F; 3H). These results establish a connection between NIK1 activation and LIMYB-mediated repression of photosynthesis-related genes, similar to the previously described control of translation-related genes mediated by the NIK1/RPL10/LIMYB signaling module.

**Figure 3.**
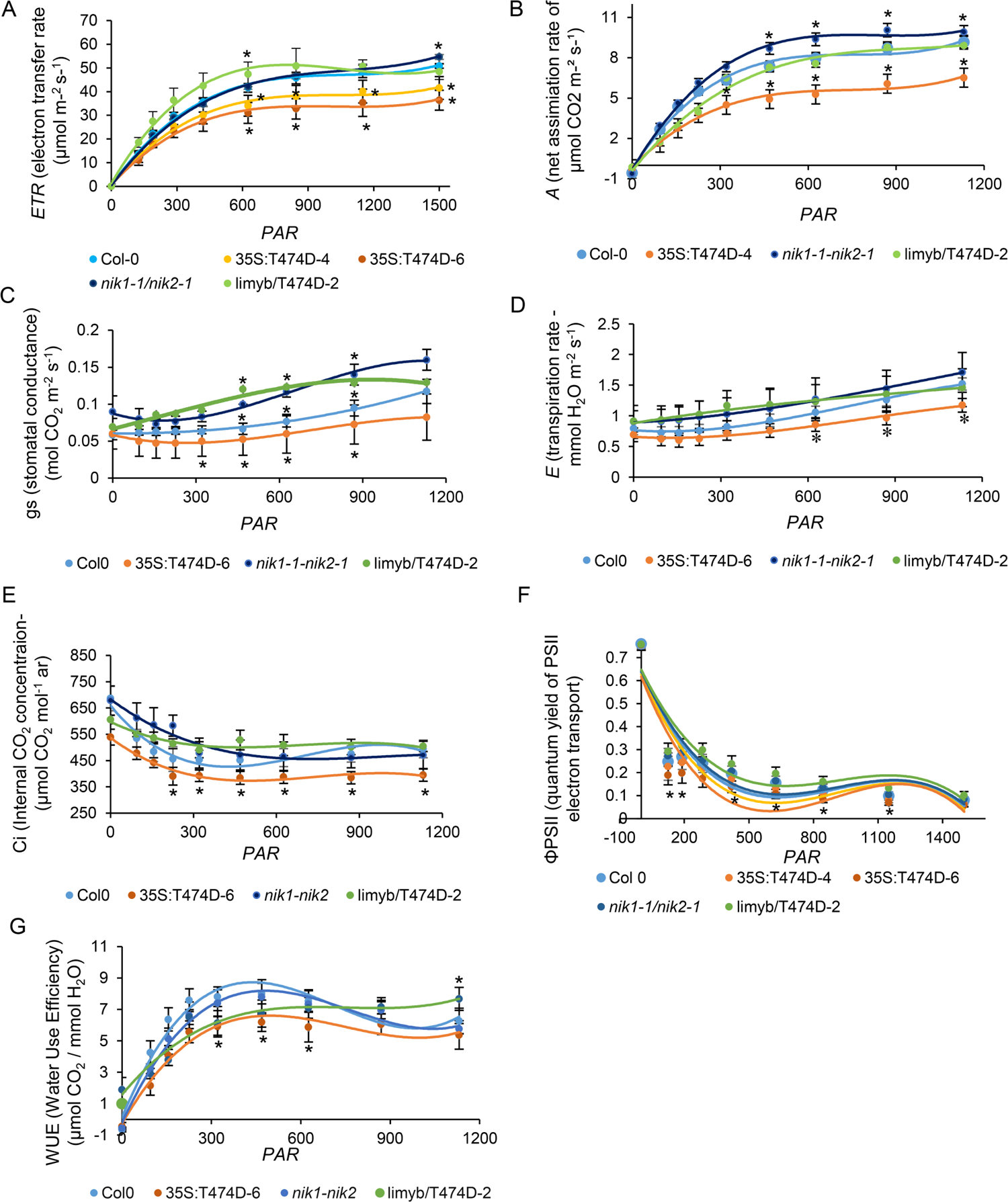
The LIMYB-mediated inhibition of photosynthesis depends on NIK1 activation. Leaf gas exchange parameters, including net CO_2_ assimilation rate (A), stomatal conductance (gs), transpiration rate (E), the internal concentration of CO_2_ (Ci), and photochemical processes, including electron transport rate (ETR), quantum efficiency, and water-use efficiency (WUE) were measured in expanded leaves from 30-days-old plants from the indicated genotypes. The experiments were repeated at least twice with similar results. Bars (±SD) with an asterisk differ from each other by the t-test (p < 0.05), n= 5.

### The promoter-repressing activity of LIMYB is regulated by NIK1-mediated phosphorylation

Although we showed that LIMYB represses the expression of photosynthesis-related genes and inhibits photosynthesis, the inactivation of *LIMYB* did not significantly alter the photosynthetic function under normal growth conditions (Figure 1E, 1F, Supplemental Figure 5). Either a functionally redundant paralog replaces LIMYB in *limyb* knockouts, or LIMYB is activated specifically under biotic stress and is therefore not required under the standard conditions of the experiment. The first hypothesis is unlikely because the closest related member of the SAINT/MYB superfamily, HAI1-Interactor 1 (HIN1), has been shown to diverge functionally from LIMYB (Chong et al., 2019). HIN1 acts as a plant-specific RNA-binding splicing regulator, invovled in osmotic stress response, interacts with serine-arginine-rich (SR) splicing factors, and increases the splicing efficiency of intron retention-prone stress-related genes. In contrast, LIMYB lacks detectable RNA binding activity (Chong et al., 2019), is dispersed evenly in the nucleoplasm (Zorzatto et al., 2015), and functions as a transcriptional repressor (Zorzatto et al., 2015; Figure 1). In addition, the HIN1 splicing function is also regulated by phosphorylation at positions S357 and S389, which are not conserved in LIMYB primary structure (Chong et al., 2019). Finally, we showed that loss-of-*LIMYB* function barely affects growth, whereas *LIMYB* overexpression inhibits growth even under normal conditions (Supplemental Figure 6), opposite phenotypes from the HUN1 transgenic lines (Cheng et al., 2019). Collectively, these data confirmed that LIMYB and HIN1 diverge functionally and may not complement each other. Therefore, we tested the second hypothesis for the weak phenotype of the *limyb* mutant under the normal experimental condition, based on the assumption that LIMYB function is activated under stress conditions, thereby not needed in the absence of biotic stimuli.

We first performed additional luciferase transactivation assays in viral PAMP-treated protoplasts to examine whether LIMYB was regulated by biotic stimuli (Supplemental Figure 9). Expression of the constitutively activated NIK1-T474D mutant repressed the RPL18 promoter activity, but not the non-target UBQ promoter, in protoplasts expressing *LIMYB* but not in non-complemented *limyb* knockouts (Supplemental Figure 9A and 9B)). These results further confirmed the specificity of LIMYB for ChiPed promoters and that NIK1-mediated downregulation of target promoters requires the LIMYB function. Monitoring LIMYB-GFP ectopic expression in *limyb* protoplasts also confirmed that the *limyb* mutant was a true knockout (Supplemental Figure 9B). Importantly, treatment of *LIMYB*-expressing protoplasts with the viral PAMP InRNA, but not UnRNA, further repressed RPL18 and PSIIOEC promoters, indicating that LIMYB may indeed be regulated by biotic stimuli. We further examined this hypothesis with complementary approaches.

As a downstream component of the NIK1 phosphorylation cascade, we examined the status of LIMYB phosphorylation under biotic stimuli known to activate NIK1 (Figure 4). Endogenous LIMYB immunoprecipitated by an anti-LIMYB antibody was probed with an anti-phosphoserine antibody after bacterial PAMP flg22 and viral PAMP InRNA treatments. Like the biotic stress-mediated phosphorylation of NIK1 and RPL10, RNA from begomovirus infected (InRNA), but not from mock-inoculated (UnRNA) Col-0 lines, induced endogenous LIMYB phosphorylation in Col-0 (Figure 4A) and YFP-LIMYB phosphorylation in the overexpressing lines L1 and L3 (Figure 4B). Likewise, the bacterial PAMP flg22 induced endogenous LIMYB (Figure 4A) and YFP-LIMYB phosphorylation (Figure 4C). Nevertheless, InRNA did not induce LIMYB phosphorylation in the *nik1/nik2* double mutant, indicating that LIMYB phosphorylation by viral PAMP required the NIK1/NIK2 function (Figure 4D). Endogenous LIMYB also required *NIK1/NIK2* function for displaying target promoter-repressing activity (Figure 4G), as InRNA treatment elicited repression of RPL18 promoter activity in Col-0 (proL18) but not in *nik1/nik2* double mutant line (proL18 + *nik1/nik2*), a phenotype that may be linked to NIK1-mediated LIMYB phosphorylation (Figure 4G).

**Figure 4.**
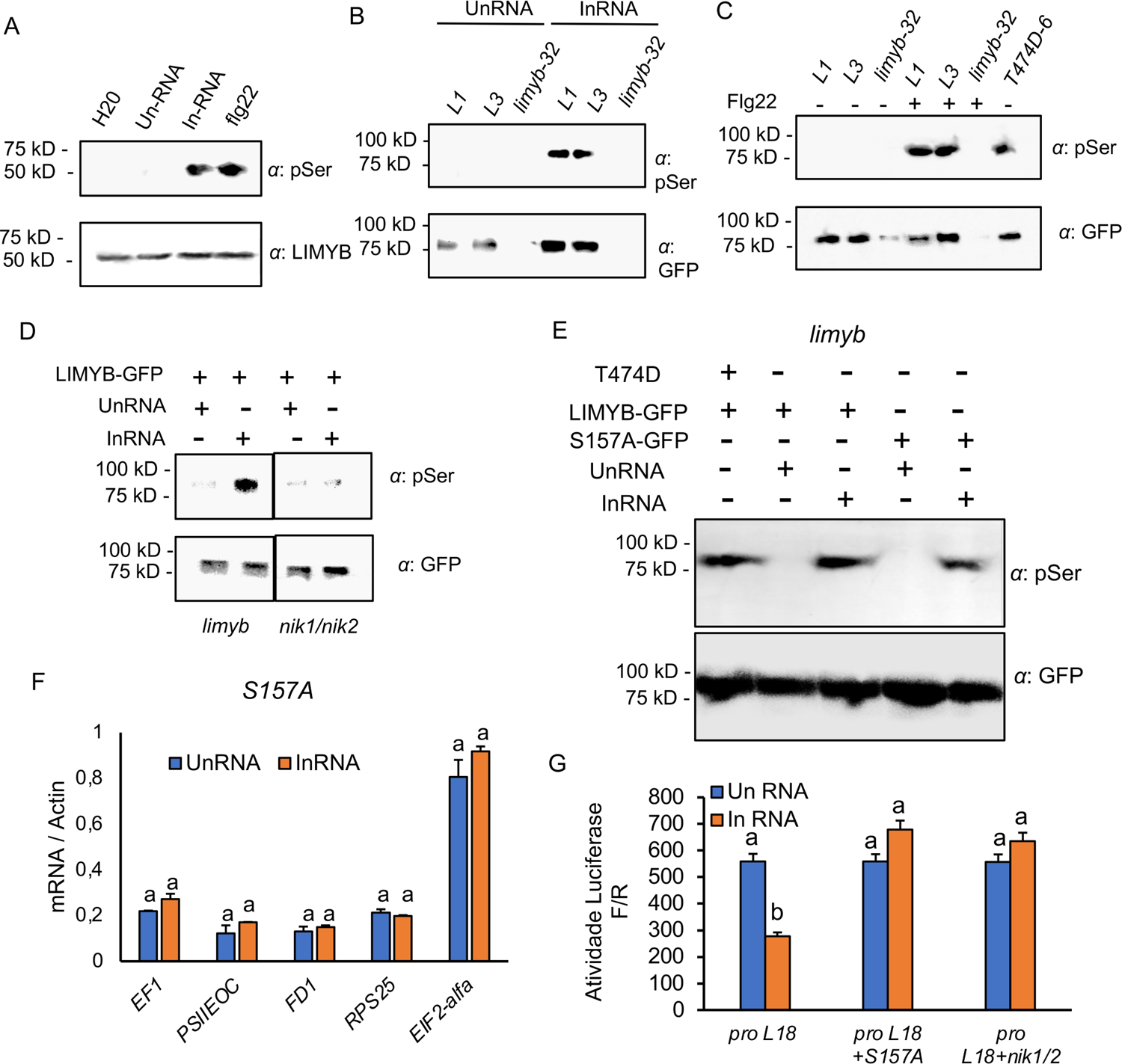
Biotic stimuli induce LIMYB phosphorylation via NIK1 activation. (A) The viral PAMP, InRN, and bacterial PAMP, flg22, induce phosphorylation of endogenous LIMYB. Col-0 seedlings were incubated with RNA prepared from begomovirus-infected leaves (InRNA), mock-inoculated leaves (UnRNA), H_2_O, and 100 mM flg22 for 3 h. LiMYB was immunoprecipitated with an anti-LIMYB polyclonal antibody and probed with an anti-phosphoserine antibody. B. UnRNA induces YFP-LIMYB phosphorylation. YFP-LIMYB-expressing Arabidopsis and *limyb* seedlings were treated with UnRNA, and InRNA for 3h. YFP-LIMYB was immunoprecipitated with an anti-GFP antibody and probed with an anti-phosphoserine antibody. C. Flg22 induces YFP-LIMYB phosphorylation. YFP-LIMYB-expressing Arabidopsis and *limyb* seedlings were treated with fgl22 for 3h. YFP-LIMYB was immunoprecipitated with an anti-GFP antibody and probed with an anti-phosphoserine antibody. NIK1-T474D-expressing line was used as a positive control for NIK1 activation. D. LIMYB phosphorylation requires the *NIK1/NIK2* function. *limyb* and *nik1/nik2* protoplasts transfected with LIMYB-GFP were treated with InRNA or UnRNA, and the immunoprecipitated LIMYB was probed with an anti-phosphoserine antibody. E. Ser157 is a possible phosphorylation site on LIMYB. The *limyb* protoplasts transfected with LIMYB-GFP or LIMYB-S157A-GFP were treated with InRNA or UnRNA and protein phosphorylation was assayed by immunoblotting with anti-phosphoserine antibody. F. UnRNA fails to cause downregulation of NIK1 signaling-associated marker genes in LIMYB-S157A-expressing *limyb* protoplasts. RNA was extracted from UnRNA and InRNA-treated protoplasts, and transcript accumulation was determined by RT-qPCR, using actin as an endogenous control for normalization. Results are mean values ±SD (n=3). Different letters denote statistically significant differences from the control line (t-test, p<0.05). G. LIMYB-S157A does not mediate the repression of the RPL13 promoter in the presence of InRNA. Col-0 protoplasts, LIMYB-S157A-expressing *limyb* protoplasts, and *nik1nik2* protoplasts were transfected with plasmids carrying the RPL13 promoter fused to luciferase and treated with UnRNA or InRNA. After 16h, luciferase activity was measured from total protein extracts of transfected protoplasts. Results are mean values ±SD (n=3). Different letters denote statistically significant differences from the control line (t-test, p<0.05).

These data are also consistent with recently published flg22-induced phosphoproteomic data, which showed that treatment with flg22 enhanced the phosphorylation signal in the LIMYB phosphosites at positions Ser157, Ser160, Ser 161, and Ser162 (fdr<0.05) (Watkins et al., 2021). Based on these data, we mutated Ser157 to Ala, creating the LIMYB-S157A mutant, and assayed for viral PAMP-induced phosphorylation. Replacing the Ser157 with Ala reduced the InRNA-induced phosphorylation in the S157A mutant, indicating that Ser157 may be phosphorylated in response to NIK1 activation (Figure 4E). The stability of the S157A mutant in protoplasts was further confirmed by immunoblotting (Supplemental Figure 9). The slight reduction of the phosphorylation signal in the S157A mutant indicated that the other flg22-induced phosphosites are also induced by a viral PAMP, keeping an almost standard phosphorylation level in the mutant. Nevertheless, InRNA did not mediate repression of translation and photosynthesis-related marker genes in *limyb* protoplast complemented with the LIMYB-S157A mutant (Figure 4F).

Furthermore, InRNA repressed the activity of the LIMYB target RPL18 promoter in Col-0 protoplasts (Figure 4G) but not in *limyb* protoplast expressing S157A. Collectively, these results demonstrated that the promoter-repressing activity of LIMYB might be regulated by PAMP-induced phosphorylation at Ser157 (Figure 4E; Watkins et al., 2021).

### NIK1 may serve as a coreceptor for different biotic stress-sensing receptors

As a member of the subfamily II of LRR-RLKs containing predominantly coreceptors of stimulus-sensing transmembrane receptors, NIK1 may be a signal relay unit from different sensing receptors (Ma et al., 2016; Hosseini et al., 2020). Accordingly, NIK1 has been shown to interact with several receptor-like kinases and may be one of the most influential information spreaders from the cell surface (Smakowska-Luzan et al., 2018; Li et al., 2019). In addition, we have previously shown that NIK1 is phosphorylated by the bacterial PAMP-induced FLS2-BAK1 immune complex activating the NIK1/RPL10/LIMYB signaling module for translation control. In contrast, we showed here that viral PAMPs can activate NIK1 and induce subsequent RPL10 phosphorylation independently of FLS2/BAK1 for NIK1, consistent with the argument that NIK1 may serve as a coreceptor from different receptors (Figure 5). In this experiment, to detect the NIK1 antiviral signaling activation, we monitored the phosphorylation of the downstream component RPL10 by probing immunoprecipitated RPL10 with an anti-phosphoserine antibody. As a positive control, we used T474D expression that induces RPL10 phosphorylation without PAMPs. While the bacterial PAMP flg22 requires the receptor FLS2 and coreceptor BAK1 to induce the NIK1-mediated phosphorylation of the downstream component RPL10 (Li et al., 2019, Figure 5A), viral PAMP induced RPL10 phosphorylation in Col-0 but also in *fls2* and *bak1* knockout lines and not in *nik1nik2* line (Figure 5B and 5C). Collectively, these results indicated that viral PAMP might require a yet-to-be-identified viral PAMP-sensing receptor for NIK1 signaling activation.

**Figure 5.**
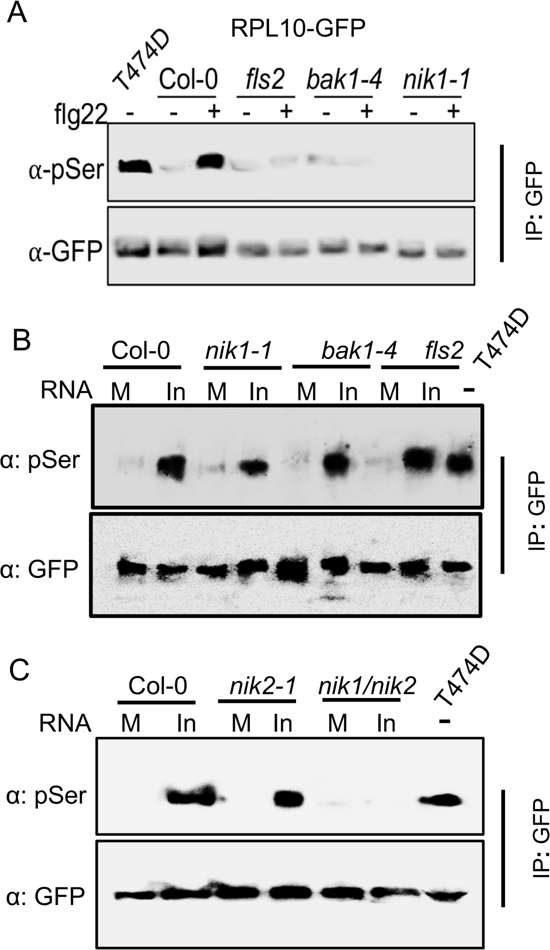
Viral PAMPs require NIK1/NIK2, but not FLS2 and BAK1, to mediate RPL10 phosphorylation. Arabidopsis protoplasts prepared from the indicated genotypes were transformed with 35S:RPL10-GFP. After 12-h for RPL10-GFP expression, the protoplasts were treated with flg22 (bacterial PAMP) **(A)** and RNA prepared from mock-inoculated (M) or begomovirus-infected (In) plants (**B, C**). Immunoprecipitated RPL10-GFP by an α-GFP antibody was probed with an α-phosphoserine antibody. T474D-6 line was used as a positive control. The experiments were repeated twice with similar results.

### The NIK1/RPL10/LIMYB signaling module is activated under abiotic stress conditions

As NIK1 may interact with different stimulus-sensing receptors, we investigated whether the NIK1/RPL10/LIMYB signaling module would function as a molecular link for the coordinate suppression of photosynthesis and translation under abiotic stress conditions. To test this hypothesis, we subjected Arabidopsis Col-0, *nik1*, *nik2*, and *nik1nik2* lines to different stress conditions and monitored the activation of the NIK1 signaling pathway. As an early response, heat promoted endogenous NIK1 phosphorylation 10 min after treatment in Col-0 and *nik2* lines but not in *nik1* and *nik1nik2* mutants, confirming they were true NIK1 knockouts (Figure 6A). As expected, T474D was constitutively phosphorylated at Ser residues. NIK1-HA also underwent phosphorylation under heat, indicating that the fusion protein was functional (Figure 6B). We also monitored the effectiveness of the heat treatment by measuring HSP70 induction (Supplemental Figure 10A). Heat treatment also promotes phosphorylation of the downstream component RPL10-GFP as early as 30 min after treatment (Figure 6C) and LIMYB (Figure 6D), followed by repression of photosynthesis-related marker genes *PHSIIOEC* and *FD1* in Col-0, but not in *nik1/nik2* knockouts (Figure 6E). Likewise, heat treatment promoted the repression of RP genes in control lines but not in the *nik1nik2* double mutant (Figure 6F). Nevertheless, loss-of-*LIMYB* function did not prevent the rapid heat stress-induced phosphorylation of NIK1 (Figure 6G) but impaired the later repression of marker gene expression in the *limyb* mutant (Figure 6H). Collectively, these data indicate that LIMYB couples the stress-induced NIK1 activation to translation and photosynthesis inhibition and may indicate that heat treatment activates the NIK1/RPL10/LIMYB signaling module to coordinate translation and photosynthesis during stress conditions.

**Figure 6.**
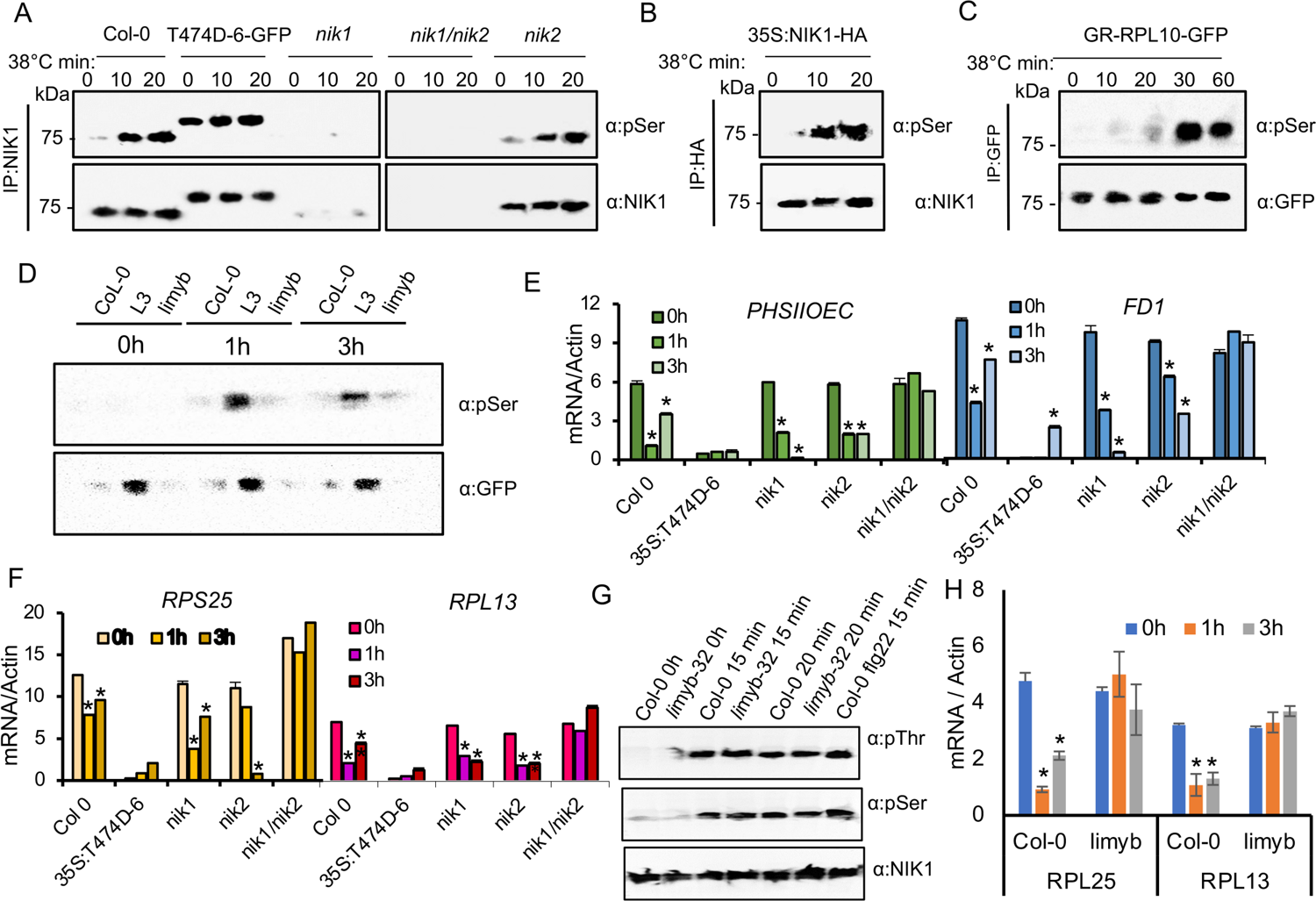
High temperature activates NIK1 antiviral signaling. **(A)** Rapid NIK1 phosphorylation in response to heat stress. Twenty-eight days-old plants were subjected to 38°C for 0 to 3h. Protein extracts were prepared at different intervals after the heat treatment. NIK1 was immunoprecipitated with an α-NIK antibody and probed with an α-phosphoserine antibody to monitor phosphorylation (top panel) and with α-NIK antibody to visualize the immunoprecipitated NIK1 (bottom panel). A T474D-expressing line was used as a positive control. **(B)** NIK1-HA is correctly phosphorylated in response to heat. NIK1-HA expressing lines were subjected to heat stress as in A. Phosphorylation was monitored as in A, except that NIK1-HA was immunoprecipitated with α-HA antibody and probed with an α-pSer antibody (top panel) and an α-NIK1 antibody (bottom panel). **(C)** Heat stress mediates RPL10 phosphorylation 30 min after the treatment. RPL10-GFP-overexpressing lines were subjected to heat treatment, and RPL10-GFP phosphorylation was monitored by probing immunoprecipitated RPL10 with an α-pSer antibody (top panel). The immunoprecipitated RPL10-GFP was also probed with an α-GFP antibody (bottom panel). **(D)** Heat stress mediates LIMYB phosphorylation. YFP-LIMYB-expressing lines were subjected to heat stress for 1h and 3h and YFP-LIMYB phosphorylation was monitored by probing immunoprecipitated LIMYB with an α-pSer antibody **(E)** Heat stress represses the photosynthetic apparatus genes PSII-OEC and FD1 in a nik1nik2-dependent manner. Plants were subjected to heat treatment for 1-h and 3-h, and the expression of the indicated genes was analyzed by qRT-PCR. Data are shown as the mean ± SE (n=3). Asterisks indicate statistically significant differences to the untreated sample (test t, P<0.05). **(F)** High temperature suppresses ribosomal protein gene expression. RP gene expression after heat stress. RT-qPCR data are shown as the mean ± SE (n=3). Asterisks indicate statistically significant differences to the untreated sample (test t, P<0.05). **(G)** Heat stress induces rapid NIK1 phosphorylation in the *limyb* knockout line. NIK1 was immunoprecipitated with an α-NIK antibody and probed with an α-phosphothreonine antibody and an α-phosphoserine antibody to monitor phosphorylation (top and middle panels) and with α-NIK antibody to visualize the immunoprecipitated NIK1 (bottom panel). **(H)** Heat stress does not induce repression of NIK1 signaling pathway-associated marker genes in *limyb*-32 mutant. The expression of the marker genes RPL25 and RPL13 was examined by RT-qPCR, 1-h, and 3-h post-treatment. Data are shown as the mean ± SE (n=3). Asterisks indicate statistically significant differences from the untreated sample (test t, P<0.05). All the above experiments were repeated at least three times with similar results.

We also examined the activation of the NIK1 signal transduction by osmotic stress, which inhibits growth (Figure 7). The effectiveness of the osmotic stress treatment was confirmed by the induction of the drought-responsive marker gene *RAB18* 3-h and 24-h after treatment in all genotypes (Supplemental Figure 10B). The osmotic stress inducer PEG mediated NIK1-HA phosphorylation (Figure 7A), leading to RPL10-GFP phosphorylation (Figure 7B), followed by repression of NIK1 activation-associated marker genes in Col-0, *nik1*, and *nik2*, but not in the *nik1nik2* double mutant (Figure 7C and 7D). Furthermore, osmotic stress induced NIK1 phosphorylation, but not the repression of the NIK1 signaling-associated marker genes, in *limyb* knockout lines (Figure 7E, 7F, and 7G), suggesting that osmotic stress-mediated NIK1 activation requires LIMYB function to repress the expression of photosynthesis and translation-related genes. Therefore, the NIK1/RPL10/LIMYB regulatory circuit is activated by different biotic and abiotic signals, further supporting a role for the NIK1 signaling pathway in coordinating translation and photosynthesis under stress conditions.

**Figure 7.**
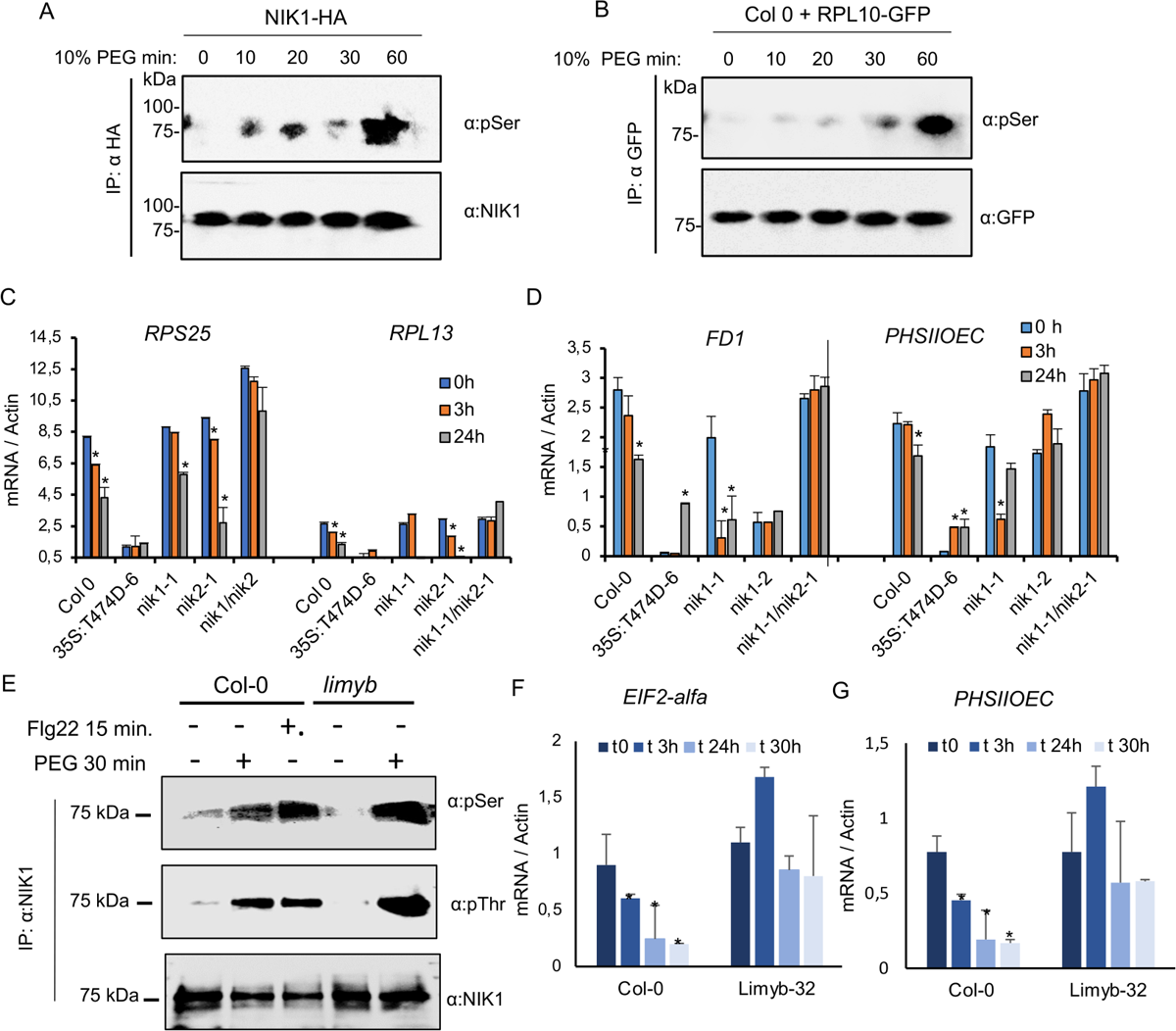
The osmotic signal triggers the NIK1 antiviral signaling activation. **(A)** NIK1 is phosphorylated under osmotic stress. NIK1-HA-expressing seedlings were treated with 10% PEG for the intervals indicated in the figure. The NIK1 phosphorylation was monitored by probing immunoprecipitated NIK1-HA with an α-pSer antibody (top panel). An α-NIK1 antibody (bottom panel) was used to visualize the immunoprecipitated NIK1-HA. **(B)** Osmotic stress induces RPL10 phosphorylation. Osmotic stress was induced in Arabidopsis seedlings expressing RPL10-GFP by PEG treatment, and then RPL10 phosphorylation was monitored by immunoblotting immunoprecipitated RPL10-GFP (bottom panel) with an α-pSer antibody (top panel). **(C)** Osmotic stress-mediated repression of ribosomal protein genes requires the function of NIK1 and/or NIK2. *RPS25* and *RPL13* expression was quantified by RT-qPCR 3-h and 24-h after PEG treatment in the genotypes indicated in the figure. A T474D-overexpressing line was used as a positive control. **(D)** Osmotic stress suppresses the photosynthetic genes *PSII-OEC* and *FD1* in Col-0 but not in *nik1nik2* double knockouts. Transcript accumulation of *PSII-OEC* and *FD1* was determined by RT-qPCR 3h and 24h after PEG treatment. **(E)** Osmotic stress induces NIK1 phosphorylation in the *limyb-32* knockout line. NIK1 was immunoprecipitated with an α-NIK antibody and probed with an α-phosphothreonine antibody and an α-phosphoserine antibody to monitor phosphorylation (top and middle panels) and with α-NIK antibody to visualize the immunoprecipitated NIK1 (bottom panel). **(F, G)** Osmotic stress does not induce repression of NIK1 signaling pathway-associated marker genes in *limyb*-32 mutant. The expression of the marker genes was examined by RT-qPCR, at the time indicated in the Figure. For **C, D, F and G,** data are shown as the mean ± SE (n=3). Asterisks indicate statistically significant differences from the untreated sample (test t, P<0.05). The above experiments were repeated at least twice with similar results.

The coordination of photosynthesis and translation allows organisms to maintain a balance between the production of chemical energy with its utilization in energy-consuming processes, particularly under stressful conditions that impact the primary metabolism of plant cells. This balance ensures efficient resource allocation and helps prevent detrimental effects such as oxidative stress. While inhibition of photosynthesis and translation negatively affects growth, as observed in transgenic lines ectopically expressing the gain-of-function NIK1-T474D mutant, any mechanism that mitigates the accumulation of oxidative stress is likely to confer a certain degree of stress tolerance. We addressed this issue by exposing the T474D lines and the loss-of-function mutants to a water deficit regime. The Col-0, *nik1-1*, *nik2-1, nik1/nik1* double mutant, *limyb-32,* and NIK1-T474D-6 lines were grown in soil for 30 days and subjected to drought by suspending irrigation until reaching the soil relative water content approximately 40%, followed by rehydration to 100% of field capacity (Supplemental Figure 11). After rehydration, the T474D lines exhibited a significantly higher survival rate compared to the wild-type line, whereas the *limyb-32* and *nik1/nik2* mutants displayed a lower survival rate than the wild-type (Supplemental Figure 11A, 11B). Throughout the water deprivation period, the T474D line maintained higher levels of leaf relative water content, while the *nik1/nik2* and *limyb-32* mutants displayed a lower water retention capacity than the wild-type (Supplemental Figure 11C). These results suggest that constitutive activation of NIK1 prevents stress build-up in response to water deprivation. This phenotype was associated with the activation of the NIK1/RPL10/LIMYB signaling module, as the loss of LIMYB and NIK1/NIK2 function compromised drought tolerance and increased susceptibility to water dehydration in the mutants.

## Discussion

All organisms must perceive and respond to changing growth conditions and environmental stimuli. During acute adverse conditions, including heat shock, osmotic stress, and pathogen attack, the induced changes in energy-consuming cellular processes must be coordinated with the rate of energy-producing metabolic processes to prevent excessive stress built up and ensure cell survival. In-plant cells, photosynthesis, a chemical energy-producing process, needs to be effectively coordinated with translation, the most energy-consuming process, under stress conditions. However, the direct connections between these two crucial cellular processes have not been thoroughly investigated. We described a growth-controlling NIK1/RPL10/LIMYB signaling module that ties the photosynthetic function to translational control in response to biotic and abiotic signals. First, we showed that LIMYB binds *in vivo* to and inhibits the activity of photosynthesis-related promoters, leading to the repression of these target genes, similar to the translation-related target genes, as demonstrated previously (Zorzatto et al., 2015). Then, we demonstrated that the LIMYB-mediated repression of the photosynthesis-related genes led to the inhibition of photosynthesis in a process that required the *NIK1/NIK2* function. We also showed that the stress-induced NIK1 phosphorylation mediated LIMYB phosphorylation, controlling the promoter-repressing activity of LIMYB in control lines but not in the *nik1/nik2* double mutant. Furthermore, the phosphomimetic, constitutively activated NIK1 mutant, NIK1-T474D, inhibited photosynthesis in control lines but not in the *limyb* mutant line. Both biotic and abiotic stimuli induced rapid NIK1 phosphorylation, followed by RPL10 and LIMYB phosphorylation, resulting in repression of the NIK1 signaling-associated marker genes in control lines but not in *limyb* knockout line. Collectively, these results demonstrated that LIMYB serves as a crucial link between biotic and abiotic-induced NIK1 activation and the repression of photosynthesis- and translation-related genes. The stress-induced LIMYB-mediated repression of these genes ensures the coordinated suppression of photosynthesis and translation. This coordination likely contributes to balance carbon fixation and allocation, precenting the occurrence of excessive oxidative stress. Therefore, the stress-induced repression of photosynthesis and translation, orchestrated by the NIK1/RPL10/LIMYB signaling module, plays a role in growth control in response to changing environmental conditions.

Compelling evidence in the literature indicates that the activation site of NIK1 signaling is the Thr-474 residue located within the kinase activation loop. This site, NIK1-T474, occupies a conserved position similar to the activation sites of BAK1 and SERK1, suggesting a similar activation mechanism among members of the LRR-RLK subfamily II (Santos et al., 2010; Yan et al., 2012). Like BAK1 and SERK1 activation sites, substituting NIK1-Thr-474 with alanine significantly reduces in vitro autophosphorylation of the NIK1 T474A kinase mutant produced in E. coli as a GST-fused protein (Santos et al., 2009). Consequently, this mutant loses its ability to promote phosphorylation and translocation of the downstream component RPL10 to the nucleus. As a result, the phosphonull mutant T474A fails to restore the antiviral function and antibacterial immunity repressing activity of NIK1 when expressed in the nik1 knockout lines, as observed in an in vivo complementation assay (Santos et al., 2009; Li et al., 2019). In contrast, previous studies have demonstrated that substituting Thr-474 with Aspartate (T474D) leads to the constitutive activation of NIK-mediated defenses in Arabidopsis and tomato plants (Zorzatto et al., 2015; Brustolini et al., 2015). Furthermore, the gain-of-function mutant T474D can sustain NIK1 phosphorylation at additional sites and induce RPL10 and LIMYB phosphorylation even without stimuli, confirming that phosphorylation of Thr-474 is the crucial event triggering kinase activation (Figure 4C and Figure 5). Supporting these data, a recent phosphoproteomic analysis discovered that Flg22 induces *in vivo* phosphorylation of NIK1/NIK2 at Thr-474, as well as Ser-465 and Ser-615 (Watkins et al., 2021). Accordingly, we have shown previously that Flg22-mediated NIK1 phosphorylation relies on BAK1 activation, which phosphorylates NIK1 at the crucial Thr-474 residue *in vitro* (Li et al., 2019).

Moreover, we have previously shown that overexpression of T474D in Arabidopsis and tomato plants results in downregulating translation-related genes and suppressing global translation in a LIMYB-dependent manner (Brustolini et al., 2015; Zorzatto et al., 2015). Similarly, our current findings demonstrate that ectopic expression of T474D induces RPL10 and LIMYB phosphorylation, repressing photosynthesis-related genes and reducing ETR and CO2 assimilation (Figures 2, 3 e 4). Therefore, any stress signal, including heat and osmotic stress, which activates the NIK1/LIMYB/RPL10 signaling circuit and downregulates translation and photosynthesis-associated marker genes is expected to induce NIK1 phosphorylation at the critical Thr-474 activation site leading to photosynthesis and translation inhibition. This stress-induced coordinated downregulation of translation and photosynthesis-related genes has been extensively demonstrated in genome-wide studies of several plant species. Consistent with our current data, analyses of previously published RNA-seq data in response to heat and osmotic stress in Arabidopsis show that prolonged stress conditions massively downregulate translation-related genes and photosynthesis-associated genes. We propose here that different stress-sensing receptors interact with and activates NIK1 relaying the stress signals to a shared signaling circuit that coordinately downregulates translation and photosynthesis. Therefore, NIK1 functions as a signaling hub for specific stress-sensing receptors.

The TOR signaling rapidly activates translation through the phosphorylation of regulatory factors in response to stimuli. In contrast, the NIK1 signaling pathway may coordinate the suppression of translation and photosynthesis during stress by regulating the repressive activity of LIMYB, which targets photosynthesis- and translation-related genes. Therefore NIK1-mediated regulation of photosynthesis and translation represents a delayed response to biotic and abiotic signals, as it depends on the half-life of the target genes. Consistent with this delayed kinetics response, the NIK1signaling pathway has been shown to suppress global translation after 8h of NIK1 activation, and photosynthesis repression may exhibit similar characteristics (Zorzatto et al., 2015). Upon NIK1 activation, LIMYB predominantly targets genes encoding components of the photosynthetic electron transport chain (ETC), photosystem II and I. Consequently, the NIK1/RPL10/LIMYB module activation leads to a simultaneous decrease in ETR, gas exchange parameters (A, gs, E, and Ci), fluorescence quantum efficiency, and WUE. These results indicate that the decline in photosynthesis activity displayed by LIMYB-overexpressing lines and T474 lines may result from the LIMYB-mediated reduction in the abundance of Photosystem II, I, and ETC components rather than from defects in C fixation or damage to the photosynthetic apparatus. Previous studies have demonstrated that adjustments in the photosystem stoichiometry can impact the quantum efficiency of photosynthesis (Chow et al., 1990).

A potential pitfall of these studies, however, is the observation that ectopic expression of the constitutively activated NIK1-T474D mutant confers tolerance to drought stress, even though the transgenic lines show reduced photosynthesis, translation, and stunted growth compared to control lines. In most cases, plants respond to low water potential by trying to avoid stress, maintaining the water potential in their tissues close to the unstressed levels. This is typically achieved by balancing the rates of water uptake and loss. In the short term, stomatal dynamics play a significant role in controlling the transpiration rate, consistent with the lower stomatal conductance observed in T474D lines. In the long term, the root-to-shoot ratio is altered, root architecture is modified to maximize water uptake, and shoot growth is reduced. There are also modifications in cuticle and lignin thickness affecting water permeability. However, avoiding a decrease in the tissue water potential under water deficit conditions comes at a cost: a reduced photosynthetic rate and reduced productivity if the stress persists. Interestingly, the T474D line displays stunted vegetative growth under normal conditions, which may have primed it for drought tolerance. Indeed, the leaf RWC of these transgenic lines under the water deficit regime remained similar to unstressed conditions, mimicking the plant’s stress avoidance mechanism to cope with drought. Additionally, the activation of the NIK1 signaling in the T474D lines may have induced mechanisms of drought tolerance. For instance, ion transporter genes, lignin biosynthetic genes, and genes involved in starvation were enriched in the set of upregulated genes by LIMYB, which requires further investigation.

Our results, along with compelling evidence from the literature, support the argument that NIK1 functions as an influential LRR-RLK information spreader and potentially acts as a coreceptor for various transmembrane stress-sensing receptors (Figure 8). Firstly, we demonstrated that NIK1 undergoes phosphorylation in response to multiple abiotic and biotic signals, activating the NIK1/RPL10/LIMYB signaling circuit, which coordinates the regulation of translation and photosynthesis in plant cells. Conceptually, signaling receptors exhibit high specificity and affinity for stimuli, making it unlikely for a single receptor to be activated by multiple signals. The versability of NIK1 in interacting with different stimulus-specific sensing LRR-RLKs, forming a hub with high centrality, supports the notion that NIK1 can transduces different stress signals as a coreceptor (Ahmed et al., 2018; Smakowska-Luzan et al., 2018; Li et al., 2019). Furthermore, NIK1 belongs to the LRRII subfamily of RLKs and shares similar structural and biochemical properties with BAK1, the almost universal coreceptor for LRR-RLKs/RLPs, (Sakamoto et al., 2012; Ma et al., 2016; Hosseini et al., 2020). Notably, the extracellular domain (ECD) of BAK1, crucial for assembling the correct receptor/coreceptor pair, is structurally equivalent to the NIK1 ECD (Jaillais et al., 2011). Finally, we have shown that NIK1 relays information from the immune complex FLS2-BAK1, transmitting biotic signals that culminate in the coordinate regulation of translation- and photosynthesis-related genes (Li et al., 2019; this work). The bacterial PAMP flg22, but not viral PAMPs, requires the immune complex FLS2-BAK1 for mediating NIK1 phosphorylation and activation. Hence, it is plausible that a yet-to-be-determined viral pattern recognition receptor might be responsible for sensing viral PAMPs and mediating NIK1 activation during viral infection. Similarly, NIK1 may transduce different abiotic signals by relaying information from other specific stress-sensing receptors. Based on these findings, we propose that NIK1 functions as a central signaling hub, integrating signals from multiple stress-sensing receptors to regulate growth under stressful conditions.

**Figure 8.**
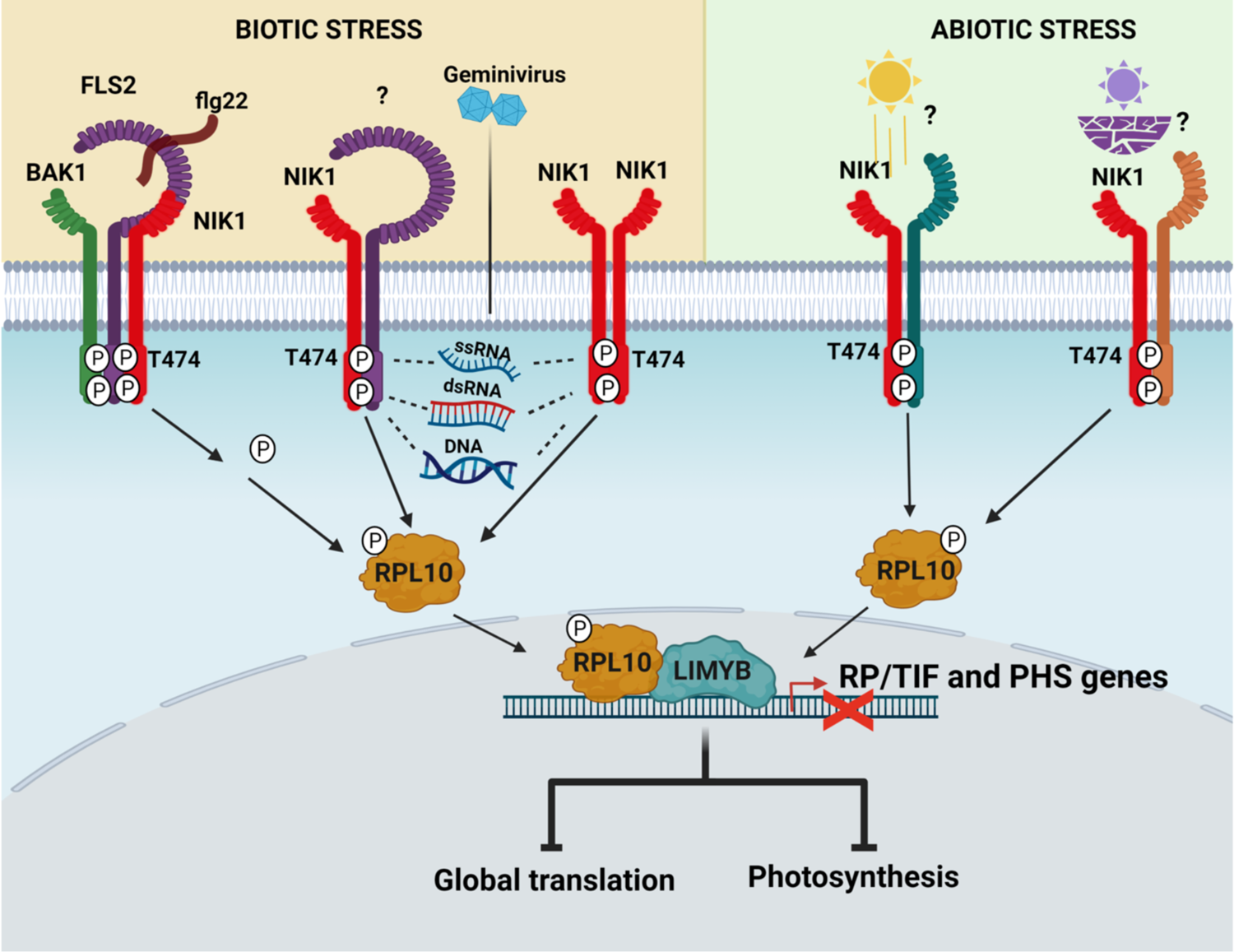
NIK1 is a signaling hub that serves as a transducer for different stress-sensing receptors. Multiple abiotic and biotic stimuli activate the NIK1/RPL10/LIMY signaling circuit, which coordinates the regulation of translation and photosynthesis in plant cells. We propose that NIK1 transduces different abiotic and biotic signals by relaying information from different stress-sensing receptors.

## METHODS

### Plant material and growth conditions

The *Arabidopsis thaliana* ecotype Columbia-0 (Col-0) was used as control lines. The knockout lines *nik1-1* (SALK_060808), *nik2-1* (SALK_044363), *fls2* (Salk_141277), *bak1-4* (Salk_116202), and *limyb-32* (SALK_032054) were obtained from the Arabidopsis Biological Resource Center and have been previously characterized (Fontes et al., 2004; Li et al., 2015; Zorzatto et al., 2015). The double mutant *nik1-1/nik2*, the transgenic line RPL10-1, harboring the construct 35S: YFP-GR-RPL10 in the Col-0 genetic background, two independently *nik1-1* transformed lines, T474D-4, T474D-6, expressing the NIK1-T474D-GFP mutant transgene, the NIK1-HA transgenic line, harboring 35S:NIK1-HA in Col-0 and the *limyb*/T474D-2 line expressing 35S:T474D-GFP in the *limyb-32* mutant background have been generated previously (Zorzatto et al., 2015; Li et al., 2019). Plants were grown in soil (Metro Mix 366) in a growth chamber at 23 °C, 45% humidity, and 75 µE/m^2^/s^1^ light with a 10-h-light/14-h-dark photoperiod. Four-week-old plants were used for treatments with viral nucleic acid elicitors, heat stress, and protoplast isolation. For osmotic stress experiments, seedlings were germinated for up to 12 days on half-strength Murashige and Skoog (MS^1^/_2_) plates containing 1% (w/v) sucrose and 0.8% (w/v) agar, grown under the same conditions.

### Plasmid construct for transient expression in protoplast

The generation of pK7F-NIK1T474D (pUFV632), which expresses a T474D mutant fused in frame with the N-terminal of GFP under the control of CaMV 35S promoter, has been previously described (Santos et al., 2009). The clones pK7F-L10 (pUFV1862) and pYFP-L10 (pUFV2365) have been previously described (Santos et al., 2009). They contain RPL10A ORF fused in frame to GFP after its last codon or YFP before its first codon under the control of the 35S promoter. The recombinant plasmid pYFP-LIMYB (pUFV1886), containing LIMYB ORF fused in-frame to the C-terminus of YFP under the control of the 35S promoter and inserted into the binary vector 35S-YFP-casseteA-Nos-pCAMBIA1300, and the recombinant plasmid AT5G05800NS-pK7FWG2 (pUVF1395), containing LIMYB ORF fused-in frame to the N-terminal of GFP under control of the 35S promoter, have been described before (Zorzatto et al., 2015).

### Plasmid constructs for promoter transactivation assay in protoplast

The clone prAt1g29970-Lucif-term-2X35S:RLucifpH7M34GW (pUVF2231), previously described (Zorzatto et al., 2015), contains the firefly luciferase cDNA under the control of the rpL18A promoter, as well as the Renilla luciferase cDNA under the control of a 2 x 35S promoter. Likewise, the clone containing the firefly luciferase cDNA under the control of ubiquitin (Ubq10) promoter for luciferase transactivation assays has been previously obtained (proUBQ:Lucif-term-2X35S:RLucifpH7M34GW; Zorzatto et al., 2015). The At2G28605 (PHSIIOEC or PsbP-OEC) promoter (−2000 bp) was obtained from *Arabidopsis* DNA by PCR using gene-specific primers (Supplemnental Table 6), and cloned into pDONR-P4-P1R vector, generating the pUFV4015 clone. The clone LUCF-term-pDON221 (pUFV2132) harbors the firefly luciferase cDNA in the entry vector pDON221, whereas the clone 2×35S-RLUCF-pDONR-P2R-P3 (pUFV2131) contains *Renilla* luciferase cDNA under the control of a 2× 35S promoter. The At2G28605 promoter (−2000 bp) was inserted into the destination vector pH7m34GW (pUFV1918), along with 2×35S-RLUCF-pDONR-P2R-P3 (pUFV2131) and LUCF-term-pDON221 (pUFV2132), by triple recombination using the gateway system (Promega, Gateway™ LR Clonase™ Enzyme mix, cat # 11791019). The resulting clone proPHSIIOEC:Lucif-term-2X35S:RLucifpH7M34GW (pUFV4027) contains the firefly luciferase cDNA under the control of the PHSIIOEC promoter (2000-bp 5’ flanking sequences), as well as the *Renilla* luciferase cDNA under the control of a 2× 35S promoter. The same procedure was used to clone the firefly luciferase cDNA under the control of the FD1 promoter (−2000 bp), yielding the clone proFD1:Lucif-term-2X35S:RLucifpH7M34GW (pUFV4031), and the truncated versions, positions −400 and −1000 bp, of proRPL18Ae promoter. The resulting clone of the truncated promoter was proL18(3):Lucif-term-2X35S:RLucif pH7M34GW (−400 bp). All promoter-specific primers used for amplifying promoter and truncated sequences are listed in Supplemental Table 6.

### Plant transformation

The *limyb-32* lines were transformed with pYFP-LIMYB using the floral dip method, generating *LIMYB-32-L1, LIMYB-32-L3,* and *LIMYB-32-L4* lines. Primary transformants were selected using 30 µgmL^-1^ hygromycin, and stable transgene insertion was monitored by PCR with gene-specific primers (Supplemental Table 6). The transgene expression level was determined by immunoblotting and real-time qPCR with gene-specific primers (Supplemental Table 6). The *actin* gene was used as an internal control to quantify gene expression by RT-qPCR.

### Whole-genome chromatin immunoprecipitation sequencing (ChIP-Seq) and data analysis

Seedlings (600 mg) harboring a YFP-LIMYB transgene were crosslinked using 1% formaldehyde in PBS solution (Sigma-Aldrich, cat. # F8775) under vacuum for 1-3 min, as previously described (Yamaguchi et al., 2014). After nuclei isolation, chromatin was sonicated up to 100–800 bp fragments. LIMYB-GFP targets from LIMYB-32-L-, LIMYB-32-L3-, and LIMYB-32-L4-isolated nuclei were immunoprecipitated using rabbit polyclonal anti-GFP - IgG antibody (Thermo Fisher Scientific, MA, USA, cat. # A11122) and captured with protein-A agarose beads (Invitrogen^TM^, Cat # 15918014). After elution, reverse crosslinking and DNA purification, ChIPed-DNA was used to generate sequencing libraries through the Illumina TruSeq kit (Illumina, cat # IP-202-1012, IP-202-1014) according to the manufacturer’s protocols. Libraries were sequenced using an Illumina HiSeq 4000. The quality of the reads was evaluated through the statistical metrics implemented in the FastQC data quality analysis tool (https://www.bioinformatics.babraham.ac.uk/projects/fastqc); low-quality sequencing reads were filtered using the printSEQ software package (Schmieder and Edwards, 2011). ChIP-Seq raw reads were aligned to the TAIR10 genome sequence using Bowtie 2 (Langmead et al., 2009). ChIP-seq peaks were calculated with the software ChIPpeakAnno (Zhu et al., 2010; Zhu, 2013), and MACS2 (Zhang et al., 2008) was used to analyze the ChIP-seq data. Putative target genes and peak distribution of the ChIP-seq data were figured out using the ChIPpeakAnno software (Zhu *et al*., 2010; Zhu, 2013). Motif discovery was performed by Multiple EM for Motif Elicitation (MEME) program (http://meme.nbcr.net/meme/), MEME-ChIP version 4.9.0 (Machanick and Bailey, 2011). Motifs were grouped by applying hierarchical clustering using motif distances, calculated by Pearson correlated coefficients as the metric for base comparison, and the Smith-Waterman Ungapped alignment method (Song et al., 2016b). The binding intensity was measured from log2 (normalized counts of reads/peaks) and evaluated at E-value cut-off of 1e-05 for sequence identification. ChIP-seq data have been deposited in the Gene Expression Omnibus under accession number GSE197332.

### RNA-sequencing (RNA-seq) method and data analysis

For RNA-seq experiments, we used three biological replicates of a pool of 12-day-old Col-0, *LIMYB-32-L1,* and *LIMYB-32-L3* seedlings and examined differences between Col-0 and LIMYB lines using the Deseq2 differential gene expression method (Robinson and Oshlack, 2010; Anders and Huber, 2010). RNA sequencing was obtained using the Illumina Hi-seq 2500 platform.

Libraries for RNA-seq were prepared with the TrueSeq-RNA Sample Prep Illumina kit (Illumina, cat # 20020595). The paired-end 100-bp protocol was used with the following quality filter parameters: 5 bases trimmed at the 3′ and 5′ ends of the reads and a minimum average Phred score of 30. Reads quality was evaluated by FastQC (https://www.bioinformatics.babraham.ac.uk/projects/fastqc). Illumina adapters were removed using Trimmomatic software (Bolger et al., 2014), and low-quality sequencing reads were filtered and trimmed using prinSEQ software package. (Schmieder and Edwards, 2011). Read mapping was performed using the Bowtie 2 program (Langmead et al., 2009). The global analysis of differential gene expression (DGE) was performed using the edger package (Langmead et al., 2009) and DEseq2 (Anders and Huber, 2010). Differential expression was determined using the p-value adjusted by the “False discover rate” (FDR). Log_2_.Fold Change p-value < 0.01 e log2-fold-change >1.0 for upregulated genes and log2-fold-change <1.0 for downregulated genes were used as the cut-off point. Differentially expressed (DE) genes were stored using SQL tables in the PostgreSQL relational database (http://inctipp.bioagro.ufv.br/, http://200.235.143.46/LiMyb/ or https://inctipp.bioagro.ufv.br/maloni/), which listed corresponding log2 FC (fold change) and p values corrected by false discovery rate (q value) for all DE genes. The classification of gene ontology (GO) and analysis of enrichment of pathways were performed using the packages GoStats (Falcon and Gentleman, 2007) and PathView (Luo and Brouwer, 2013), both present in the R/Bioconductor GOstats repository. The significant p-value for enrichment was P < 0.01.

### Activation of the NIK1 pathway by begomovirus-derived nucleic acids and flg22

Leaf discs from transgenic lines previously injured with an abrasive were excised by a cork borer and incubated overnight in 12-well plates containing 500 µL ultra-pure water. Then the water was immediately replaced with 500 µL TE buffer supplemented with 250 ng/µL of viral nucleic acids (DNA or total RNA prepared from begomovirus-infected plants and treated with proteases, Rnase, or DNase), 100 mM flg22 or ultra-pure water. NIK1 activation was monitored by phosphorylation at 10 to 60 min after treatment. Phosphorylation of RPL10 was examined at 30 min to 3 h post-treatment, whereas the transcript accumulation of target genes was measured at a minimum of 3 h after treatment.

### Protoplast preparation from *A. thaliana* leaves

Expanded leaves from 3–4-week-old plants were cut into 0.5–1-mm pieces and digested with 5– 10 ml enzyme solution (20 mM MES pH 5.7; 1.5% cellulase R10; 0.4% macerozyme R10; 0.4 M mannitol and 20 mM KCl). The leaf strips were vacuum infiltrated for 30 min in the dark using a desiccator. Then, the digestion was conducted without shaking in the dark for at least 3-h at room temperature. The integrity of the protoplasts was examined under a light microscope. The protoplasts were filtered to remove undigested leaf tissues and recovered by centrifugation. The protoplasts were transformed using the protocol of DNA-PEG–calcium transfection, in which 10 μl DNA (10–20 μg of plasmid DNA) were mixed with 2 × 10^4^ protoplasts along with 5% PEG solution (Polyethylene glycol 4000, sigma, Cat # 25322-68-3) at room temperature for up to 15 min and then incubated for recovery during 12 h (Yoo and Sheen, 2007).

### Heterologous expression of 6xHis-NBSNIK1 and 6His-LIMYB and production of anti-NIK1 and anti-LIMYB antibodies

The 6xHis-NBSNIK1 recombinant plasmid has been previously cloned in pDEST17 via LR-mediated recombination (Gateway Invitrogen Life Technologies, Inc). It harbors the NIK1 kinase domain fused to 6xHis under the control of an isopropyl-β-d-thiogalactopyranoside (IPTG) inducible T7 promoter. Likewise, the 6xHis-LIMYB recombinant plasmid was generated by transferring LIMYB ORF from pDON-R201 (Zorzatto et al., 2015) to the pDEST17 vector via LR-mediated recombination (Gateway Invitrogen Life Technologies, Inc). An aliquot of each plasmid DNA was used to transform E. coli strain C41(DE3). For the heterologous expression, a pre-inoculum was diluted 1:100 in 100 mL of LB medium, and the culture was incubated at 37°C, under agitation at 225 rpm until reaching optical density (OD600nm) between 0.6 and 0.8. The recombinant protein synthesis was induced by 0,8 mM IPTG for 4 h at 37 °C. Cells were harvested, washed, and suspended in lysis buffer (50 mM Tris pH 8.0, 50 mM NaCl, 5mM EDTA, 100 µg/ml lysozyme, 0.1% Triton X-100, and 1 mM PMSF) before use. Cell pellets were suspended in 1/10 of the original volume of the culture and incubated at 30 °C for 5 minutes. Lysis was obtained by ultrasonication in 6 cycles of 10 seconds on/10 seconds off, amplitude 20%, keeping the tube on ice throughout the procedure. Finally, cell debris was removed by ultracentrifugation at 4 °C for 20 min at 14000 g. The His-tagged fusions were affinity purified using Ni-NTA agarose (Quiagen®), according to the manufacturer’s recommendations, and resolved by SDS-PAGE.

For antibody production, two male BALB/c mice (4- to 8-week-old) were used. The animals were obtained from the Central Animal Laboratory of the Center of Biosciences and Health of the Universidade Federal de Viçosa (Brazil) (CEUA/UFV – process number 45/2013). The mice were maintained and handled in the experimental vivarium of the Immunology and Virology sector of the General Biology Department – DBG/UFV, where they remained in 12 h light/dark photoperiod cycle and received water and food ad libitum. Immunization was performed intraperitoneally with 200 µL of a solution prepared from 3 to 4 protein bands, cut directly from the polyacrylamide gel, and ground with a tissue homogenizer in PBS buffer added with 50 µg of saponin. The treatment consisted of 3 doses each 15 days. The blood was collected from the animals, and the serum was heated in a thermoblock at 56 °C for 30 minutes to inactivate the complement proteins and then titrated by dot blot.

### Phosphorylation assay

Protoplasts of *nik1-1, nik2-1, fls2, bak1-4,* and *nik1-1/nik2-1* mutant lines were transformed with RPL10-GFP. After 16-h incubation for transient transgene expression, the protoplasts were treated with nucleic acids prepared from mock-inoculated and begomovirus-infected plants or 100 mM flg22 for 3-h. RPL10-GFP was immunoprecipitated with α-GFP antibodies and protein-A agarose beads, fractionated by SDS-PAGE, and immunoblotted with an α-phosphoserine antibody (α-phosphoserine peroxidase, Sigma-Aldrich, Cat # SAB5200087, 1:5000) and an α-GFP antibody (α-GFP, life technologies, cat # A11122, 1:5000; goat anti-mouse IgG-HRP, Santa Cruz, Cat # sc-2005, 1:10,000). Likewise, RPL10-GFP- and NIK1-HA-expressing transgenic lines were treated for 3-h with nucleic acids prepared from uninfected and begomovirus-infected plants or 100 mM flg22. Then, RPL10-GFP and NIK1-HA were immunoprecipitated from leaf discs from treated and non-treated plants with an α-GFP antibody or α-HA antibodies (α-HA, Thermo Fisher, Cat # 71-5500, 1:50) and protein-A agarose beads (InvitrogenTM, Cat # 15918014), fractionated by SDS-PAGE and immunoblotted with an α-phosphoserine antibody (α-phosphoserine peroxidase, Sigma-Aldrich, Cat # SAB5200087, 1:5000) or α-phosphothreonine antibody (α-phosphothreonine, Thermo Fisher, Cat # 71-8200, 1:250) and goat anti-rabbit IgG secondary antibody (Thermo Fisher, Cat # 65-6120, 1:10.000). The same protocol examined the heat and osmotic-induced phosphorylation of NIK1-HA and RPL10 in 22-day-old transgenic lines.

### RNA isolation and RT-qPCR analysis

For RNA isolation, either 12-day-old seedlings grown on half-strength MS plates or leaf discs of 22-day-old plants were transferred to 1 mL H_2_O in a 12-well plate to recover overnight and then treated with uninfected and begomovirus-infected nucleic acid preparation or 100 nM flg22 for 15 min to 3h, or 10% PEG 8000 for 3-h and 24-h. RNA was extracted using TRIzol reagent (Invitrogen^TM^, cat # 15596018, USA) and quantified with a NanoSpec spectrophotometer. Total RNA was treated with DNase I, RNase-free (Thermo Scientific™, cat # EN052) for 30 min at 37 °C and then reversed transcribed with M-MLV Reverse Transcriptase (Invitrogen™, cat # 28025013). Real-time PCR was performed using iTaq Universal SYBR Green Supermix (Bio-Rad, cat # 1725121, USA) and a 7500 Fast Real-Time PCR System (Applied Biosystems™, cat # 4351106, USA). The expression of each gene was normalized to the expression of Actin or UBQ10. The primers used to analyze specific transcripts for RT-qPCR are listed in Supplemental Table 6.

### Promoter transactivation assay in protoplasts and *Nicotiana benthamiana* leaves

The protoplasts were isolated from Col-0 and *limyb-32*, and co-transformed with the combinations: pK7F-NIK1T474D + proL18(2):Lucif-term-2X35S:RLucifpH7M34GW (−2000 bp); pK7F-NIK1T474D + AT5G05800NS-pK7FWG2 + proL18(2):Lucif-term-2X35S:RLucifpH7M34GW; AT5G05800NS-pK7FWG2 + proL18(2):Lucif-term-2X35S:RLucifpH7M34GW; pK7F-NIK1T474D + proUBQ:Lucif-term-2X35S:RLucifpH7M34GW; pK7F-NIK1T474D + AT5G05800NS-pK7FWG2 + proUBQ:Lucif-term-2X35S:RLucifpH7M34GW. *N. benthamiana* leaves were agroinfiltrated with *A. tumefaciens* GV3101 strains carrying the following combinations in the absence and presence of LIMYB cDNA (AT5G05800NS-pK7FWG2): proPHSIIOEC:Lucif-term-2X35S:RLucifpH7M34GW (−2000 bp), proFD1:Lucif-term-2X35S:RLucif pH7M34GW (−2000 bp); proL18Ae(1):Lucif-term-2×35S:RLucif pH7M34GW (−1000 bp); proL18(3):Lucif-term-2X35S:RLucif pH7M34GW (−400 bp), proUBQ:Lucif-term-2X35S:RLucifpH7M34GW. Forty-eight hours after infiltration, 200 mg of leaf tissue were harvested for total protein extraction. Luciferase activity was assayed with Dual-Luciferase® Reporter Assay System (Promega, cat # E1910) according to the manufacturer’s instructions.

### Photosynthetic and photochemical analysis of LIMYB and T474D overexpressing lines

Gas exchange and chlorophyll fluorescence measurements were carried out in expanded leaves from 30-day-old LIMYB and T474D lines, grown in a chamber at 23°C. Gas exchange rates were determined in the morning, applying different photon rates m^-2^s^-1^ for photosynthetic efficiency analysis, using an infrared gas concentration determination system (IRGA, LCpro-SD - ®ADC Bio scientific, Hoddesdon, England). The net assimilation of carbon (A, µmol CO_2_ m −2 s −1), stomatal conductance (gs, mol H_2_O m-2s −1), the internal concentration of CO_2_ (Ci, µmol CO_2_ mol-1), transpiration rate (E, mmol m-2 s-1) and water-use efficiency (WUE, A/E μmol m-2 s-1/mmol H2O m-2 s-1), under CO_2_ concentrations controlled by the use of a gas cylinder attached to the equipment. Photochemical parameters of PSII and the energy transmission of the light-capturing complex (Lchb) were determined with a modulated light fluorometer (MINI-PAM, Walz, Effeltrich, Germany). Minimum (Fo) and maximum (Fm) fluorescence of dark-adapted leaves were also determined in the morning to analyze FSII Effective quantum yield (YII) and electron transport rate (ETR = ΦFSII x RFA x 0,5 x 0,84) (Genty et al., 1989). ETR rate was determined via modulated light pulse, the response curve of chlorophyll-a fluorescence parameters (ETR, ΦFSII, ΦNO, and qL) in responses to eight increasing light levels based on Fo and Fm values (minimum value up to the saturating pulse).

### Heat stress

The heat stress treatment was performed as described (Chao et al., 2017; Wang et al., 2020), with some modifications. Twenty-eight-day-old plants were subjected to 38°C for 0 to 3-h. NIK1 phosphorylation was monitored 0, 10, 15, 20, 30, and 60 min post-treatment. As for RPL10 phosphorylation and transcript accumulation of target genes, 0, 1-h, and 3-h were taken after treatment. After the specified treatment period, the leaves were immediately frozen in liquid nitrogen for protein and/or total RNA extraction.

### Osmotic stress

Osmotic stress was induced in Arabidopsis seedlings as described (Costa et al., 2008), with some modifications. 12-days-old *Arabidopsis thaliana* seedlings were germinated on Murashige and Skoog (MS) medium containing 1% sucrose (pH 5.7) in a petri dish and then placed in 12-well plates containing 90% liquid MS medium (1/4 strength) and 10% polyethylene glycol (PEG, MW 8000, SIGMA, cat # 25322-68-3) (p/v) for 3- and 24-h.

### Measurement of growth and developmental parameters

Development and growth rate of Col-0, T474D-4, T474D-6, *nik1-1, nik2-1, nik1-1/nik2-1*, LIMYB-32-1, LIMYB-32-3, and *limyb-32*, were monitored as previously described (Chirinos-Arias and Spaminato, 2020), with some modifications. Germination was evaluated from seed protrusion in medium MS^1^/_2_. Primary root growth was measured daily for seedlings grown in vertically positioned plates. Vegetative growth was studied in soil-grown plants subjected to short diurnal periods, photoperiod 8h-light/16h-dark, or long 10h-light/14h-dark period; the digital tool Image J (http://rsb.info.nih.gov/ij/) was used to analyze rosette area and total leaf number. The transition from the vegetative to the adult phase was examined from the emergence of the floral bud, days to the first flower opening, and the number of primary and lateral inflorescences. The mean number of siliques and seeds per silique were also analyzed. The number of siliques was determined in primary and lateral shoots; seed number per silique was recorded from siliques taken from the main shoot positioned between 6 and 12 from the initial silique. The total number of seeds was estimated as the number of seeds per silique x the number of siliques.

### Statistical analysis

The Student’s t-test was used for statistical analysis, performed in Excel 2019 software. The analyses were executed with three-independent experiments, and the data were expressed as means ± sd. A significant difference was considered p-value ≤ 0.05. The photosynthetic parameters analysis used 3° order polynomial regression on adjusted radiation curves and Student’s t-test to compare curvature points.

## AUTHOR CONTRIBUTIONS

RMT, MAF, and EPBF conceived the project, designed experiments, and analyzed data. OJBB performed the bioinformatics analysis and the statistical analysis of the data. TFFS, JJ-B, SSB, INS, NGAR, CEMD, and LLL performed experiments and analyzed data. LLO, HJOR, and PABR analyzed data and provided critical feedback. RMT, MAF, and EPFB wrote the manuscript with inputs from all co-authors.

## ACKNOWLEDGMENTS

This research was supported by Conselho Nacional de Desenvolvimento Científico e Tecnológico (CNPq Grant no. 441955/2019-3 and 403819/2021-0 to E.P.B.F.) and Fundação de Amparo à Pesquisa do Estado de Minas Gerais, Brazil (Fapemig Grants no APQ-01282-17 and RED-00205-22 to E.P.B.F.). R.M.T. was supported by graduate fellowships from Fapemig and CNPq, respectively. T.F.F.S., J.J.-B., and S.S.B were recipients of CAPES graduate fellowships. M.A.F. and C.E.M.D. were pos-doctoral fellows from CNPq and Capes, respectively. I.N.S. and N.G.A.R. were recipients of undergraduate scholarships from CNPq.

## DATA AND MATERIALS AVAILABILITY

Data are available in the main text or the Supplementary Material. ChIP-seq data have been deposited in the Gene Expression Omnibus under accession number GSE197332.

