## Supplemental Figures S1-S11 for "Coordinated regulation of photosynthesis and translation via NIK1/RPL10/LIMYB signaling module in response to biotic and abiotic stresses"

### Supplementary Figures

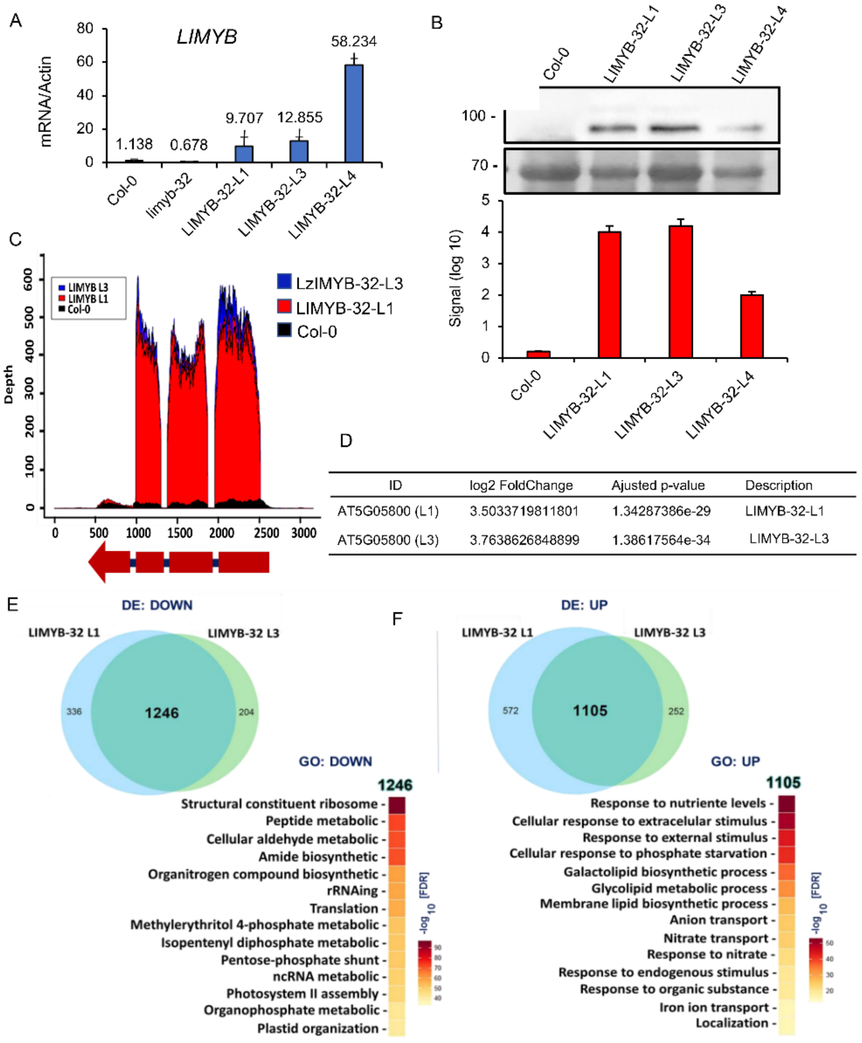


**Supplementary Figure 1. Induced transcriptome by LIMYB overexpression.** (**A)** Accumulation of YFP-LIMYB transcripts in the transgenic lines LIMYB-32-L1, L3, and L4. Gene expression was analyzed by RT-qPCR. Absolute expression was calculated using the $2^{-\Delta Ct}$ method and actin as an endogenous control. The bars represent 95% confidence interval based on replicates from 3 independent experiment samples. **(B)** Accumulation of recombinant protein YFP-LIMYB in LIMYB-32-L1, LIMYB-32-L3, and LIMYB-32-L4. Total protein extracted from the transgenic lines was separated by electrophoresis, transferred to membranes, and probed with polyclonal α-GFP antibody. Bands were quantified using the Quantity One BioRad program. (**C)** Schematic representation of the LIMYB Locus (At5g05800) with mapped RNA sequencing hits. The gene contains three introns and four exons. The relative abundance of RNA hits at the At5g05800 locus is shown in black for Col-0, red for L1, and blue for L3. (**D)** LIMYB relative expression level in LIMYB-32-L1 and LIMYB-32-L3 expressing lines compared to Col-0 as determined by RNA-seq. **(E)** Venn diagram depicting the overlap between down-regulated genes in LIMYB-32-L1 and LIMYB-32-L3. Significantly enriched GO terms (false discovery rate, FDR of - log 10) derived from genes that were downregulated in both LIMYB-32-L1 and LIMYB-32-L3 lines compared to Col-0. (**F)** Venn diagram indicating the overlap between upregulated genes in LIMYB-32-L1 and LIMYB-32-L3. Heat map of gene ontology (GO), representing the distribution of functional categories of commonly upregulated genes; DE analyses of GO used an FDR value of - log 10.

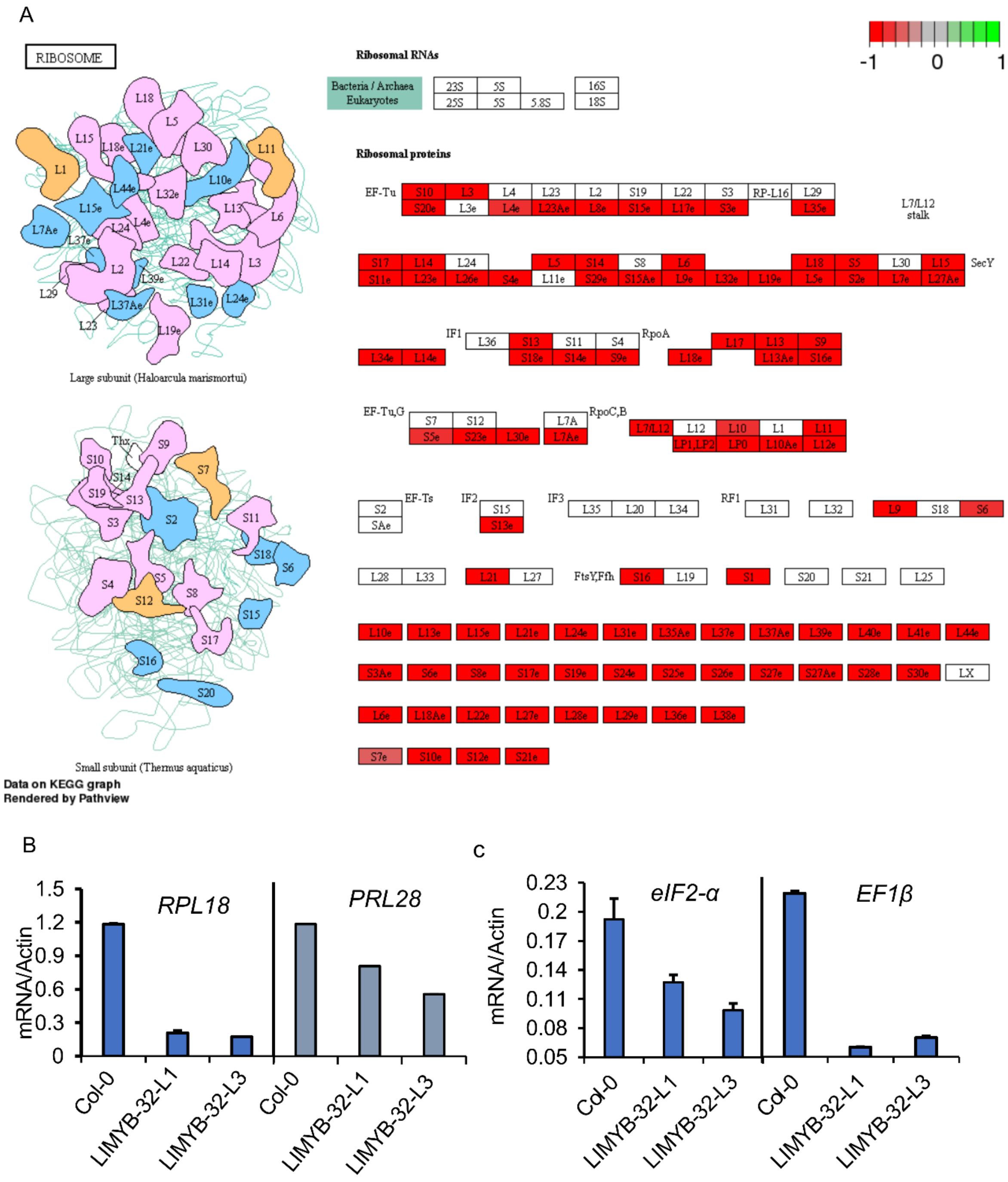


**Supplementary Figure 2. LIMYB overexpression down-regulates translation-related genes.** (**A**) KEGG illustration of the differential expression analysis. The significant predominance of the ribosomal protein gene network in the LIMYB-mediated downregulated changes. (**B**), (**C**) LIMYB overexpression represses the expression of ribosomal protein genes and translation initiation and elongation factor-encoding genes. Gene expression was determined by RT–qPCR of mRNA from LIMYB-overexpressing and Col-0 seedling. Values represent the mean ± SD of five biological replicates.

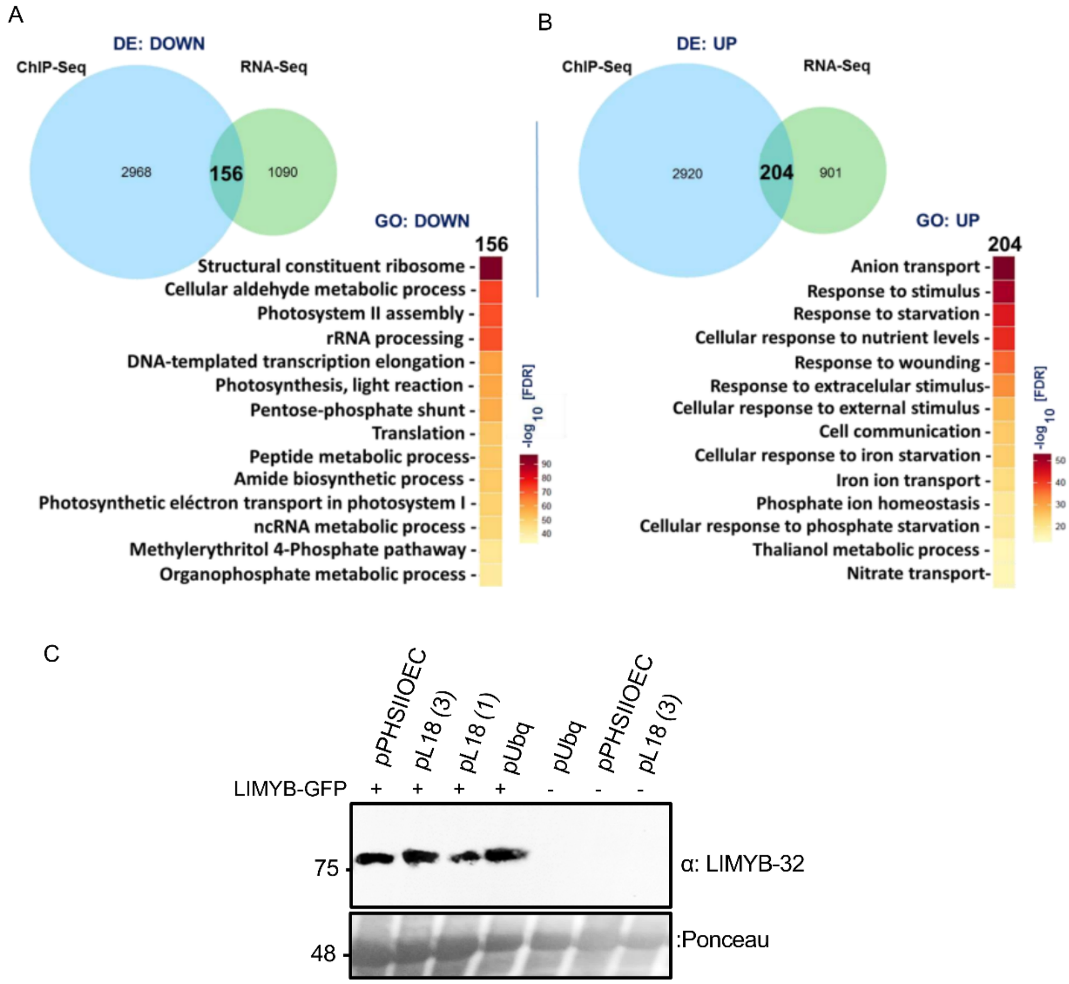


D

ChIPSeq/RNASeq

| Id gene | Upstream | log2FC | Log10 q-value | padj |
|---|---|---|---|---|
| proL18Ae | - 455 | - 0.62 | 28.30 | 5.5e-15 |
| proPHSIIOEC | - 1574 | -1.67 | 15.14 | 4.1e-19 |

**Supplementary Figure 3. Integrative analysis between ChIP-seq and RNA-seq on LIMYB-overexpressing lines defines the LIMYB regulon.** (**A**) Venn diagram indicating the overlap between down-regulated genes by LIMYB (RNA-seq) and LIMYB target genes (ChIP-seq). Heat map of significantly enriched GO terms (false discovery rate, FDR of - log 10) derived from the LIMYB down-regulon. (**B**) Venn diagram indicating the overlap between upregulated genes by LIMYB (RNA-seq) and LIMYB target genes (ChIP-seq). Heat map of significantly enriched GO terms (false discovery rate, FDR of - log 10) derived from the LIMYB up-regulon. **(C)** *LIMYB* expression in *N. benthamian* leaves for the promoter transactivation assay. LIMYB protein expression was detected by immunoblotting protein extracts from agroinfiltrated leaves using an α-GFP antibody. **(D)** Upstream position of LIMYB motifs and level expression detected by ChIP-seq and RNA-seq, respectively.

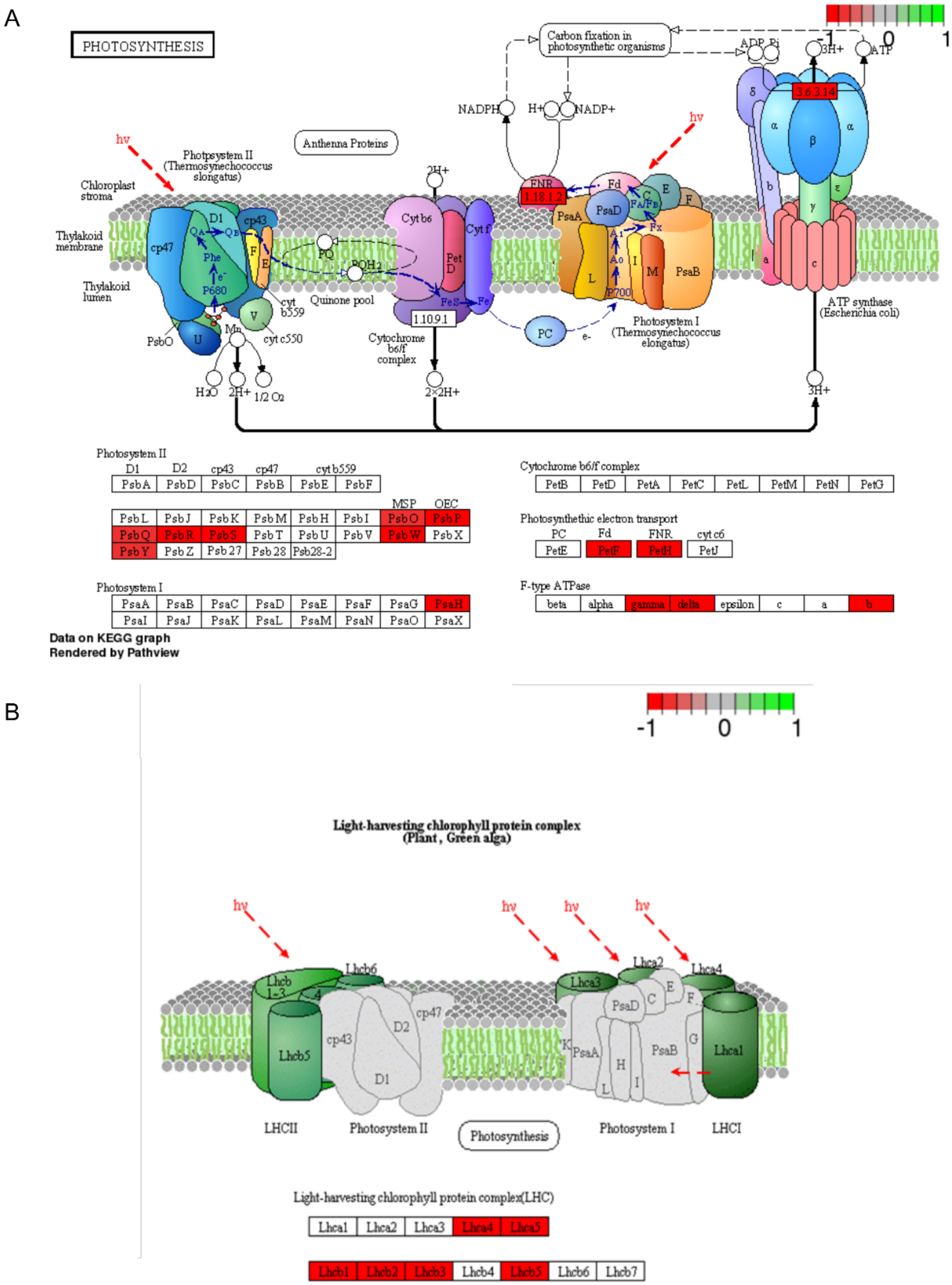


**Supplementary Figure 4. Representation of photosynthetic apparatus-related genes in the LIMYB down-regulon. (A)** Significant repression of subunits of PSII, PSI, and ATP synthase. (**B)** Expression of light-collecting complex (LHC I and LHC II). Differential expression analysis demonstrates that LIMYB regulates the expression of proteins related to the assembly of photosynthetic complexes.

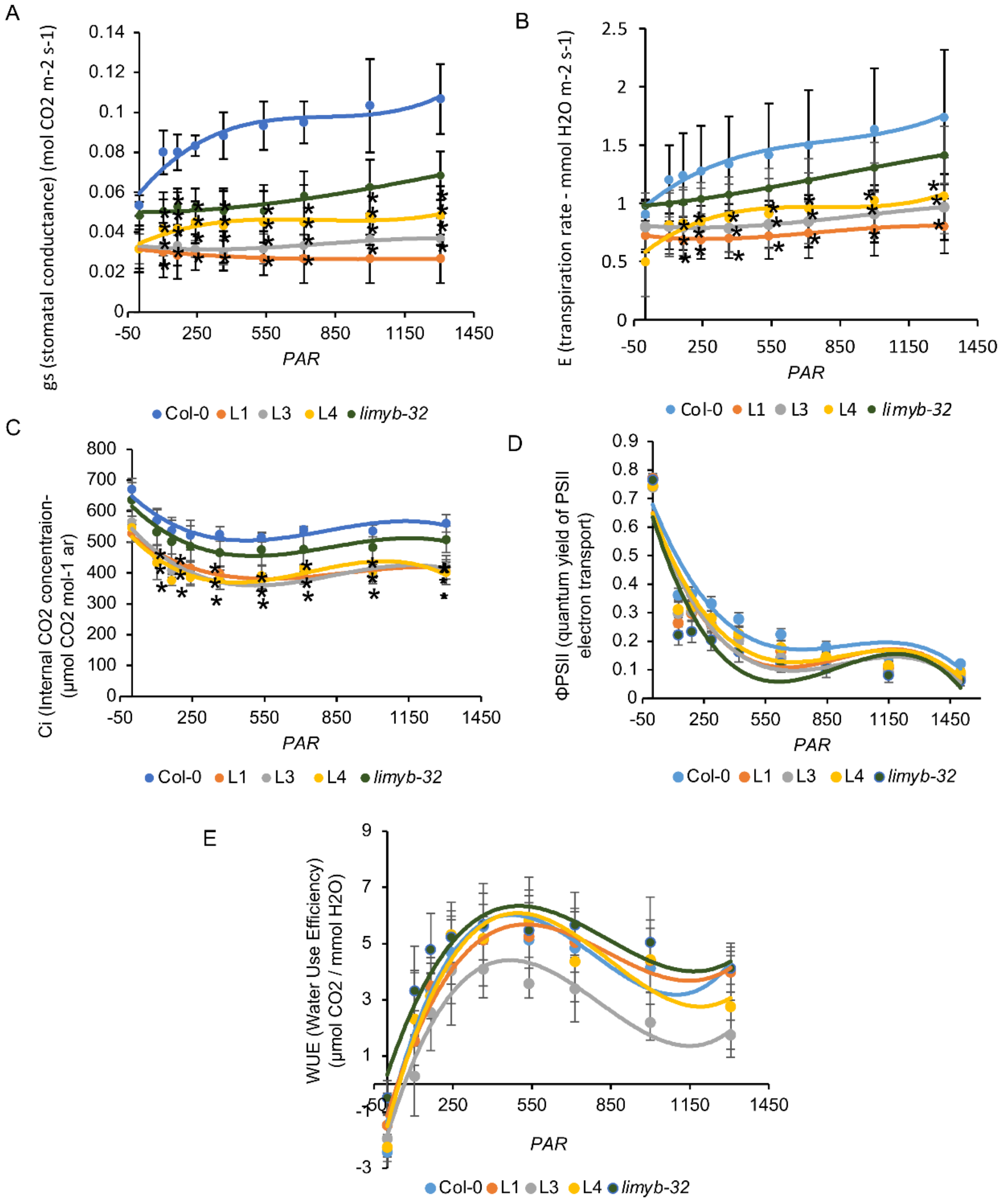


**Supplementary Figure 5. LIMYB affects the photosynthetic activity in plants. (A)** Stomatal conductance (gs), **(B)** transpiration rate (E), **(C)** the internal concentration of $CO_2$ (Ci), **(D)** the quantum efficiency of yield (ΦPSII), **(E)** water use efficiency (WUE), of LIMYB-overexpressing lines and Col-0. Data are mean values ±SD (n=6). Asterisks indicate significant differences from the control by the Student's t-test ($p < 0.05$). Col-0, Columbia. L1, LIMYB-32-L1. L3, LIMYB-32-L3. L4, LIMYB-32-L4, and *limyb*, knockout line. The above experiments were repeated twice with similar results

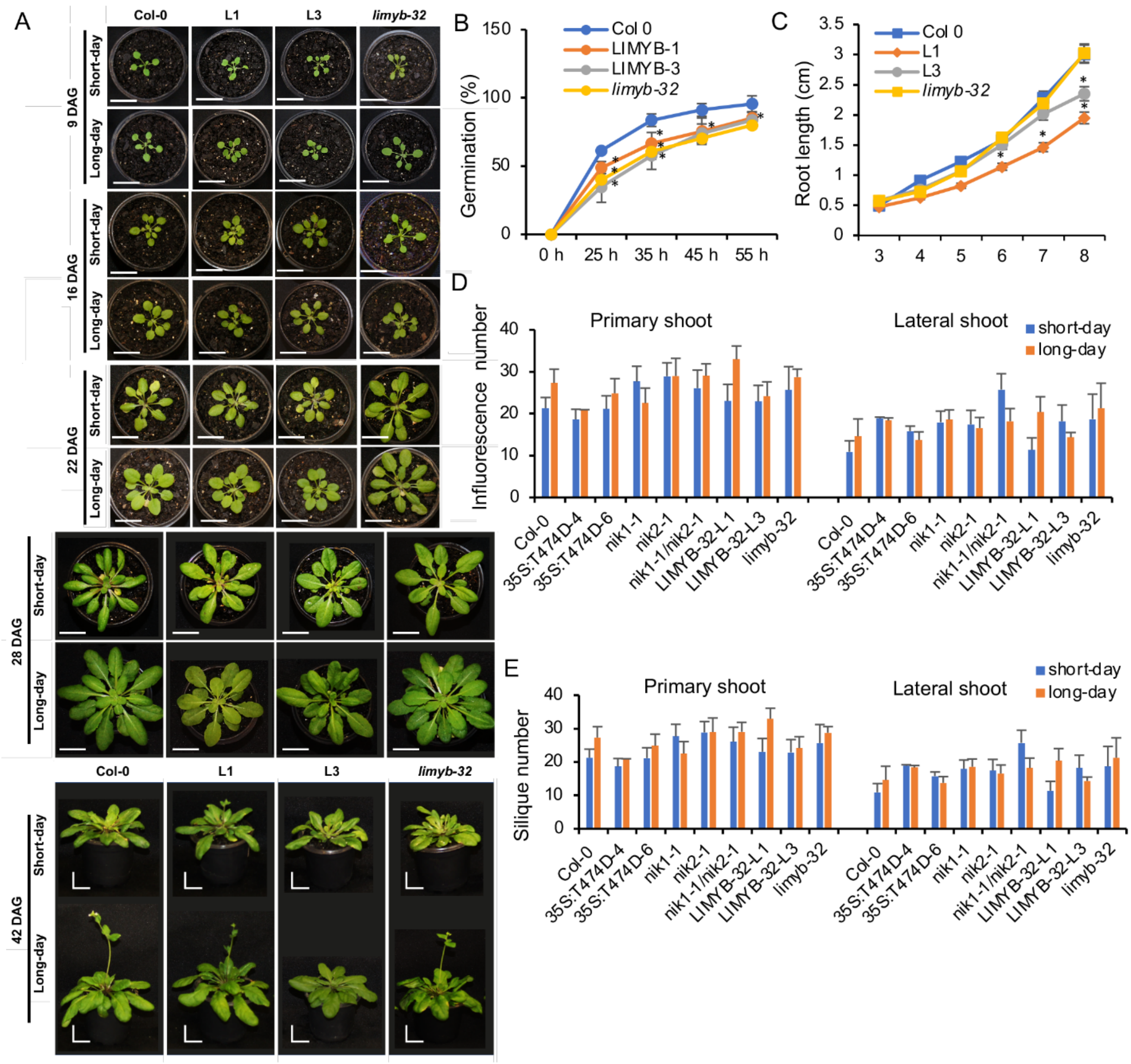


**Supplementary Figure 6. LIMYB overexpression affects growth and development**. **(A)** The growth phenotypes of transgenic lines during the vegetative and reproductive phases. Growth phenotypes were recorded 9 to 48 days after germination under short-day and long-day photoperiods. **(B)** Seed germination, **(C)** root length, **(D)** inflorescences number, **(E)** silique number were determined for LIMYB-overexpressing lines and *limyb* knockout line in comparison with Col-0. Data are mean values ±SD (n=6). Asterisks denote significant differences from the control by the Student's t-test ($p < 0.05$).

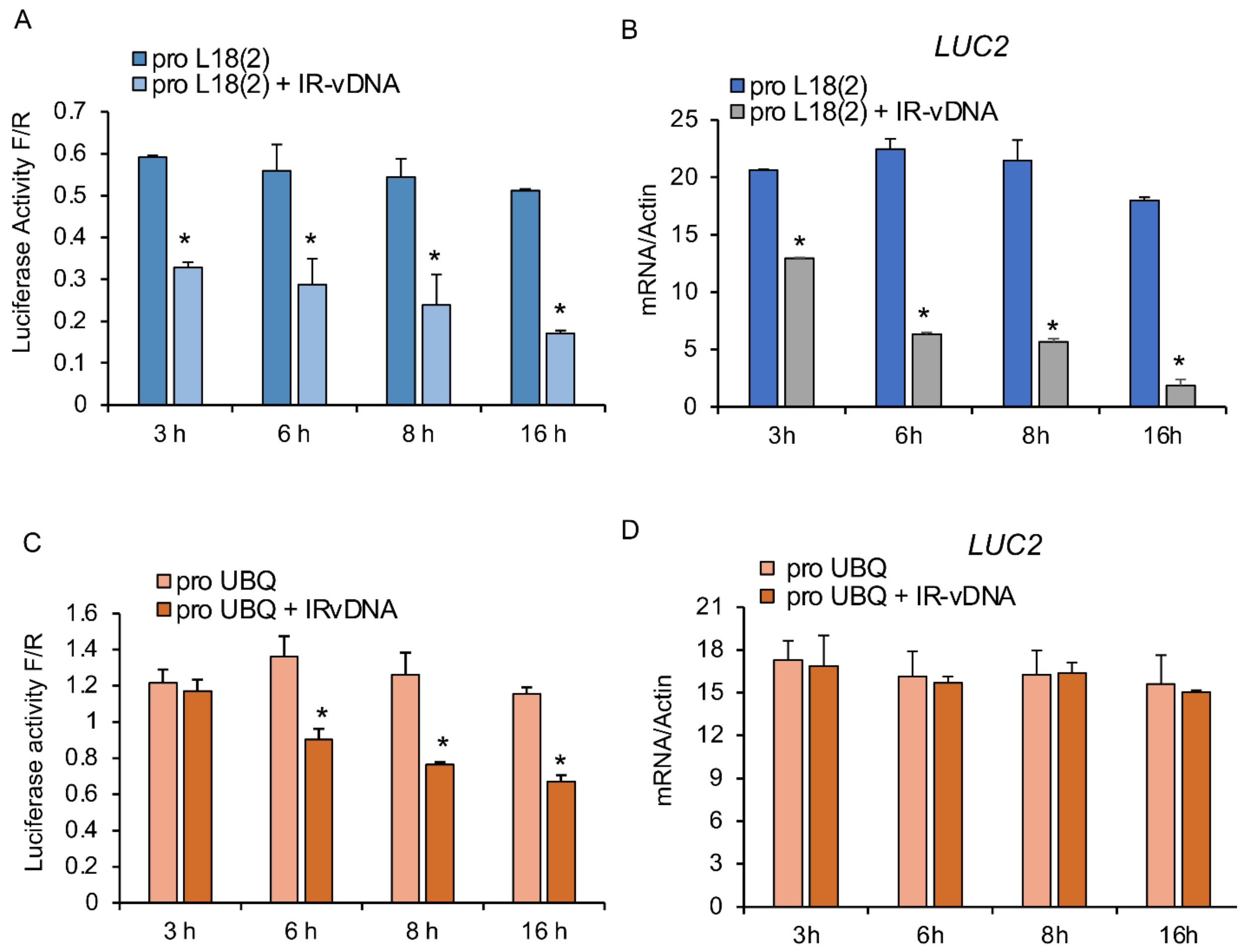


**Supplementary Figure 7. Protein synthesis inhibition by LIMYB is a late response.** Protoplasts co-transformed with 35S:LIMYB-GFP + proL18(2) –Luciferase (proL18(2) ) or proUBQ-Luciferase (proUBQ) were treated with a viral PAMP (IR-vDNA; intergenic region from the component B of Begomovirus, and luciferase activity and mRNA accumulation were measured in the intervals of 3, 6, 8 and 16 h. **(A), (B)** Time course of LIMYB-controlled luciferase activity and mRNA accumulation under the control of RPL18 promoter. F/R, firefly luciferase activity to Renilla activity ratio. **(C), (D)** LIMYB does not target the UBQ promoter but represses luciferase activity as a late response. Progression time of luciferase activity and mRNA accumulation of the UBQ promoter after LIMYB expression. Three independent replicates were used ($n = 3$). Asterisks indicate statistically significant differences from the control by the test-t ($p < 0.05$).

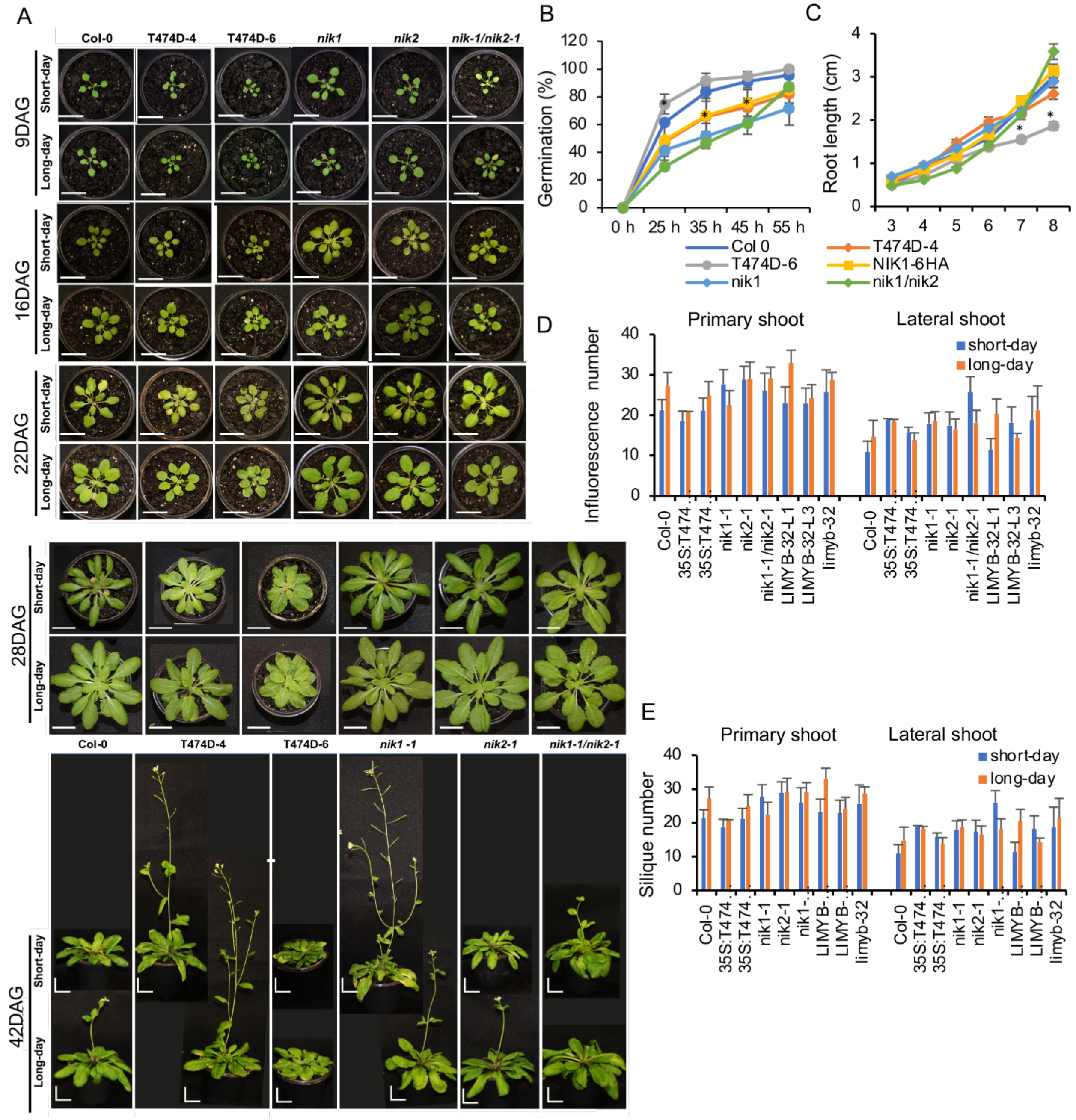


**Supplementary Figure 8. Activation of NIK1 antiviral signaling affects growth and development. (A)** The growth phenotypes of T474D-expressing lines, *nik1*, *nik2*, and *nik1nik2* knockout lines and Col-0 during the vegetative and reproductive phases were recorded from 9 to 48 days after germination under short-day and long-day photoperiods**. (B)** Seed germination, **(C)** root length, **(D)** inflorescences number, **(E)** silique number of T474D-expressing lines and Col-0 were determined. Bars (± SD, n=6) with an asterisk differ from each other by the t test ($p < 0.05$), n = 2.

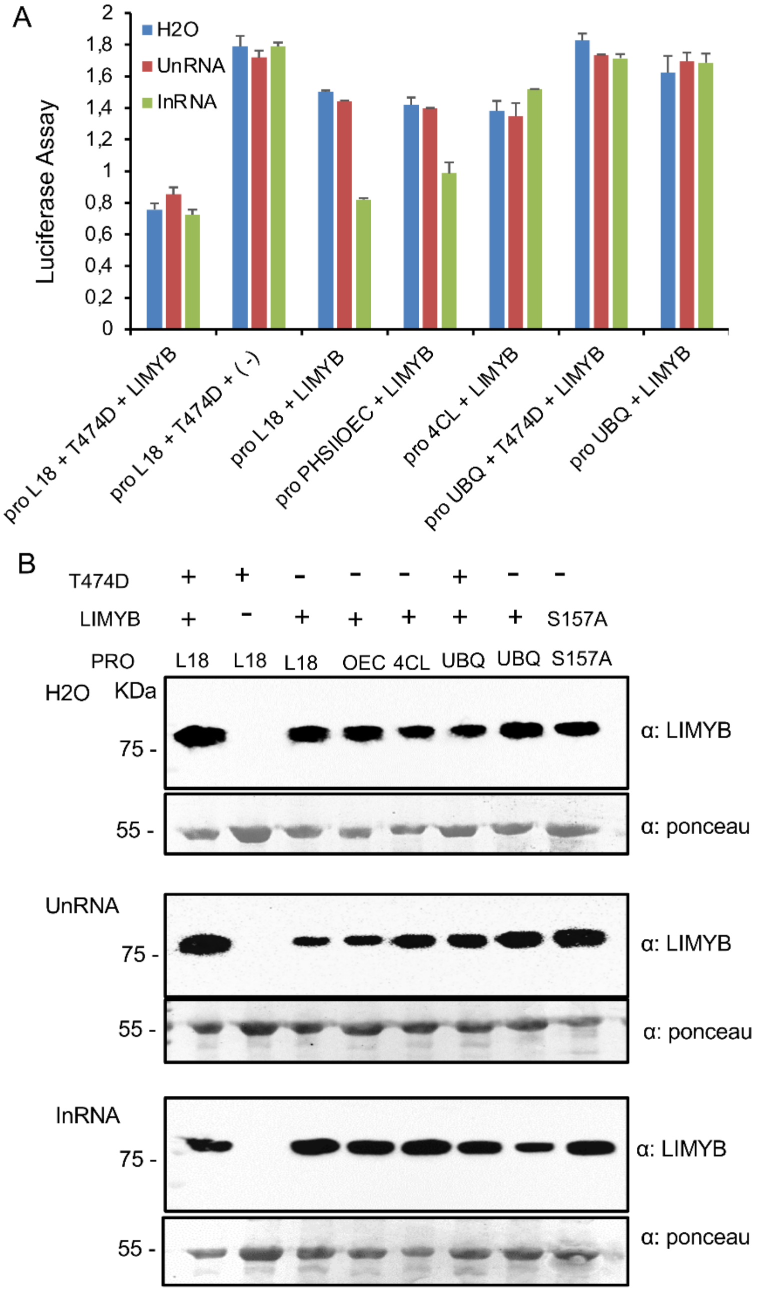


**Supplementary Figure 9. InRNA enhances further the promoter-repressing activity of LIMYB**. **(A)** The *limyb* protoplast was transfected with plasmids carrying the indicated promoters fused to luciferase in combination with T474D or LIMYB-expressing constructs, as indicated in the figure, and treated with $H_20$, UnRNA or InRNA. After 16h, luciferase activity was measured from total protein extracts of transfected protoplasts. Results are mean values ±SD (n=3). B. Immunoblotting of LIMYB in transfected protoplasts. Total protein was fractionated by SDS-PAGE and immunoblotted with an anti-LIMYB antibody. The last lane, S157A, corresponds to the accumulation of the S157A-GFP mutant in *limyb* protoplasts transfected with S157A-GFP.

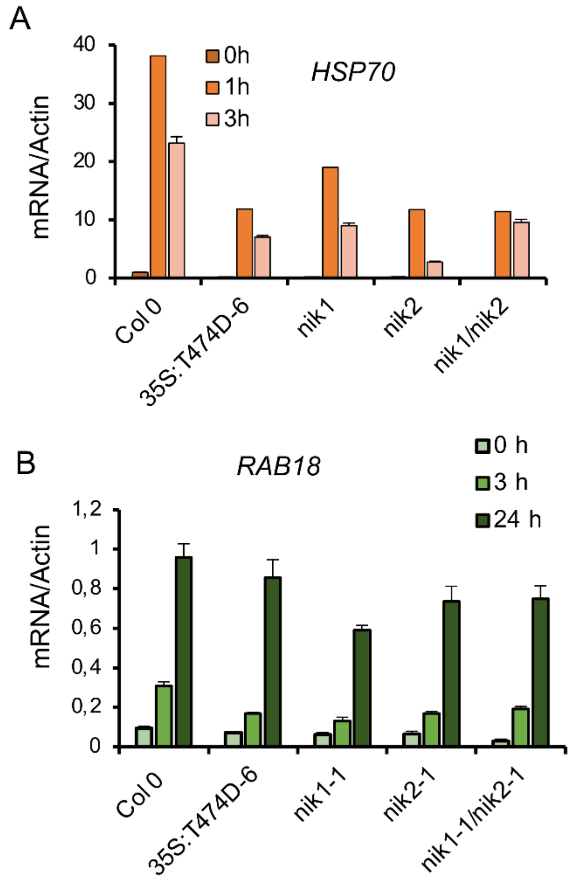


**Supplementary Figure 10. Transcript accumulation of heat stress- and osmotic stress-associated marker genes. (A)** The heat stress was effective for all genotypes examined. The induction of the heat-associated marker gene, HSP70, was examined by RT-qPCR after 1h and 3h of the treatment. **(B)** Osmotic stress induction by PEG treatment in all genotypes examined. The expression of the osmotic stress-associated marker gene RAB18 was examined by RT-qPCR 3-h and 24-h post-treatment.

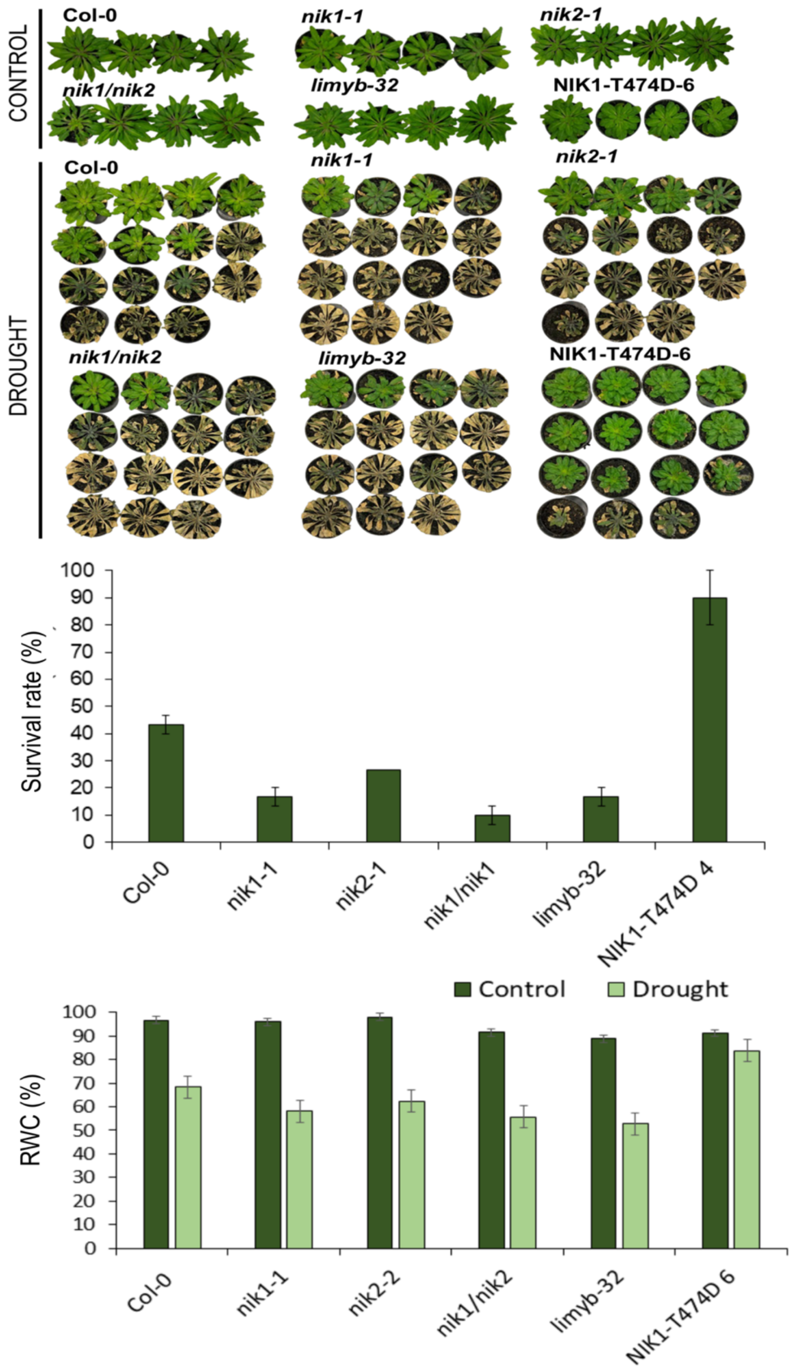


**Supplementary Figure 11. NIK1 signaling induces drought tolerance in Arabidopsis. (A)** T474D is more tolerant to drought. The Col-0, *nik1-1*, *nik2-1, nik1/nik1* double mutant, *limyb-32* and NIK1-T474D-6 lines were grown for 30 days in soil and subjected to drought by suspending irrigation until reaching the RWC approximately 40%, followed by rehydration to 100% of field capacity. The top part of the panel represents control plants grown under normal conditions. **(B)** Survival rate of Col-0, *nik1-1*, *nik2-1, nik1/nik1,* limyb-32 and NIK1-T474D-6 lines after rehydration. The values represent the average (± *SD)* of three independent experiments with a total of 45 individuals evaluated for drought resistance. **(C)** Relative water content of (RWC) 3 days before rehydration. Values represent the average (± *SD)* of 5 evaluated individuals.
