## Supplemental Tables S1-S6 for "Coordinated regulation of photosynthesis and translation via NIK1/RPL10/LIMYB signaling module in response to biotic and abiotic stresses"

**Supplemental Table 1.** Categories downregulated by LIMYB overexpression, present in the integrative RNA-seq/ChIP-seq analysis.

| Down regulated GO_ID | Pvalue | OddsRatio | ExpCount | Count | Size | Term |
|---|---|---|---|---|---|---|
| GO:0003735 | 0.000 | 17.270 | 22 | 187 | 398 | structural constituent of ribosome |
| GO:0006081 | 0.000 | 6.955 | 19 | 95 | 339 | cellular aldehyde metabolic process |
| GO:0010207 | 0.000 | 6.512 | 2 | 12 | 177 | photosystem II assembly |
| GO:0006364 | 0.000 | 6.907 | 3 | 18 | 256 | rRNA processing |
| GO:0006354 | 0.000 | 4.981 | 7 | 30 | 133 | DNA-templated transcription, elongation |
| GO:0019684 | 0.002 | 4.293 | 2 | 7 | 156 | photosynthesis, light reaction |
| GO:0006098 | 0.000 | 4.673 | 2 | 10 | 200 | pentose-phosphate shunt |
| GO:0006412 | 0.000 | 4.761 | 6 | 25 | 526 | translation |
| GO:0006518 | 0.000 | 5,298 | 7 | 31 | 566 | peptide metabolic process |
| GO:0043604 | 0.000 | 5 | 7 | 30 | 578 | amide biosynthetic process |
| GO:0009773 | 0.000 | 22.606 | 3 | 29 | 51 | photosynthetic electron transport in PHS I |
| GO:0034660 | 0.000 | 6.546 | 23 | 110 | 413 | ncRNA metabolic process |
| GO:0019288 | 0.005 | 3.186 | 3 | 8 | 229 | methylerythritol 4-phosphate pathway |
| GO:0046490 | 0.006 | 3.129 | 3 | 8 | 233 | isopentenyl diphosphate metabolic process |
| GO:0090407 | 0.003 | 1.684 | 24 | 38 | 446 | organophosphate biosynthetic process |

**Supplemental Table 2.** Categories upregulated by LIMYB overexpression, present in integrative RNA-seq/ChIP-seq analysis.

| Up regulated GO_ID | Pvalue | OddsRatio | ExpCount | Count | Size | Term |
|---|---|---|---|---|---|---|
| GO:0006820 | 0.000 | 4.271 | 32 | 110 | 619 | anion transport |
| GO:0050896 | 0.000 | 1.895 | 74 | 126 | 1814 | response to stimulus |
| GO:0042594 | 0.000 | 9.613 | 16 | 100 | 306 | response to starvation |
| GO:0031669 | 0.000 | 9.878 | 16 | 103 | 310 | cellular response to nutrient levels |
| GO:0009611 | 0.000 | 2.845 | 17 | 44 | 335 | response to wounding |
| GO:0009991 | 0.000 | 8.089 | 19 | 105 | 363 | response to extracellular stimulus |
| GO:0071496 | 0.000 | 8.795 | 18 | 107 | 349 | cellular response to external stimulus |
| GO:0007154 | 0.000 | 6.558 | 21 | 102 | 439 | cell communication |
| GO:0010106 | 0.000 | 6851 | 6 | 31 | 116 | cellular response to iron ion starvation |
| GO:0006826 | 0.000 | 5.413 | 1 | 6 | 99 | iron ion transport |
| GO:0030643 | 0.000 | 3.412 | 5 | 7 | 362 | phosphate ion homeostasis |
| GO:0016036 | 0.000 | 2.657 | 3 | 7 | 124 | cellular response to phosphatte starvation |
| GO:0080003 | 0.000 | 5.897 | 2 | 3 | 3 | thalianol metabolic process |
| GO:0010167 | 0.000 | 6.295 | 10 | 49 | 197 | response to nitrate |
| GO:0015706 | 0.000 | 6.564 | 11 | 53 | 207 | nitrate transport |

**Supplemental Table 3.** Ribosomal protein and translation-associated genes downregulated by LIMYB overexpression, present in integrative RNA-seq/ChIP-seq analysis.

| Gene_id | width | Fold enrich. | log10 q-value | Distance to feature | log2 FC | padj | Description |
|---|---|---|---|---|---|---|---|
| AT1G09690 | 185 | 4.13 | 10.52 | -152 | -0.77 | 0.00 | Translation protein SH3-like family protein |
| AT1G22780 | 173 | 4.74 | 13.92 | -867 | -1.65 | 0.00 | Ribosomal protein S13/S18 family |
| AT1G74060 | 371 | 6.04 | 23.03 | -79 | -0.33 | 0.03 | Ribosomal protein L6 family protein |
| AT2G27530 | 184 | 5.23 | 16.85 | -144 | -1.40 | 0.00 | Ribosomal protein L1p/L10e family E |
| AT2G34480 | 222 | 6.93 | 28.30 | -1334 | -0.55 | 0.00 | Ribosomal protein L18ae/LX family protein |
| AT2G36620 | 184 | 4.62 | 13.22 | -1250 | -1.26 | 0.00 | ribosomal protein L24 RPL24A |
| AT2G37270 | 271 | 7.79 | 36.16 | -75 | -0.69 | 0.00 | ribosomal protein 5B |
| AT3G07110 | 172 | 5.11 | 16.10 | -350 | -1.16 | 0.00 | Ribosomal protein L13 family protein |
| AT3G09200 | 188 | 4.27 | 15.26 | -75 | -0.66 | 0.00 | Ribosomal protein L10 family protein |
| AT3G25520 | 173 | 5.84 | 20.74 | -455 | -1.18 | 0.00 | ribosomal protein L5 |
| AT3G27830 | 212 | 6.69 | 26.56 | -220 | -1.15 | 0.00 | ribosomal protein L12-A |
| AT3G44590 | 279 | 9.70 | 54.85 | -1091 | -1.69 | 0.00 | 60S acidic ribosomal protein family |
| AT3G53890 | 201 | 5.71 | 19.94 | -340 | -1.68 | 0.00 | Ribosomal protein S21e |
| AT3G55280 | 184 | 5.18 | 19.86 | -1065 | -0.72 | 0.00 | 60S ribosomal protein L23A (RPL23aB) |
| AT4G13170 | 237 | 7.05 | 29.18 | -595 | -1.56 | 0.00 | Ribosomal protein L13 family protein |
| AT4G31985 | 193 | 6.80 | 29.21 | -2389 | -1.46 | 0.00 | Ribosomal protein L39 family protein |
| AT4G35490 | 180 | 5.96 | 21.54 | -302 | -0.67 | 0.00 | mitochondrial ribosomal protein L11 |
| AT4G39200 | 230 | 5.59 | 19.15 | -793 | -1.22 | 0.00 | Ribosomal protein S25 family protein |
| AT5G02870 | 264 | 9.24 | 46.22 | -402 | -0.50 | 0.01 | Ribosomal protein L4/L1 family |
| AT5G13510 | 230 | 8.66 | 54.93 | -630 | -1.08 | 0.00 | Ribosomal protein L10 family protein |
| AT5G22440 | 235 | 10.45 | 60.19 | -693 | -1.34 | 0.00 | Ribosomal protein L1p/L10e family |
| AT5G27770 | 303 | 4.28 | 11.84 | -364 | -1.49 | 0.00 | Ribosomal L22e protein family |
| AT5G46160 | 205 | 8.51 | 40.31 | -972 | -1.07 | 0.00 | Ribosomal protein L14p/L23e family |
| AT5G59850 | 266 | 11.19 | 62.97 | -624 | -1.44 | 0.00 | Ribosomal protein S8 family protein |
| AT5G64140 | 170 | 5.35 | 17.61 | -222 | -0.99 | 0.00 | ribosomal protein S28 (RPS28); |
| AT1G09690 | 185 | 4.13 | 10.52 | -152 | -0.77 | 0.00 | Translation protein SH3-like family protein |
| AT1G22780 | 173 | 4.74 | 13.92 | -867 | -1.65 | 0.00 | Ribosomal protein S13/S18 family |
| AT1G74060 | 371 | 6.04 | 23.03 | -79 | -0.33 | 0.03 | Ribosomal protein L6 family protein |
| AT2G27530 | 184 | 5.23 | 16.85 | -144 | -1.40 | 0.00 | Ribosomal protein L1p/L10e family |
| AT2G34480 | 222 | 6.93 | 28.30 | -1334 | -0.55 | 0.00 | Ribosomal protein L18ae/LX family protein |
| AT2G36620 | 184 | 4.62 | 13.22 | -1250 | -1.26 | 0.00 | ribosomal protein L24. cytosolic |
| AT2G37270 | 271 | 7.79 | 36.16 | -75 | -0.69 | 0.00 | ribosomal protein 5B |

**Supplemental Table 4.** Translation Factors downregulated by ectopic expression of LIMYB.

| **Gene_id** | **width** | **Fold enrich.** | **log10 q-value** | **Distance to feature** | **log2 FC** | **padj** | **Description** |
|---|---|---|---|---|---|---|---|
| AT4G11175 | 216 | 5.471 | 18,375 | -2400 | -1.57 | 0 | Translation initiation fator IF-1 (Chloroplastic) |
| AT5G12110 | 185 | 6.32 | 24.02 | -363 | -1.16 | 0 | Translation elongation factor EF1B |
| AT2G36010 | 357 | 9.76858 | 52.78026 | -1750 | -0.98 | 0 | Eukaryotic translation initiation factor 2 α |
| AT5G05470 | 244 | 7.17 | 30.07 | -863 | -0.9 | 0 | eukaryotic translation initiation factor 3 |
| AT1G70782 | 252 | 8.27491 | 40.50072 | -829 | -0.82 | 0 | E2F transcription factor 3, sub. G. |
| AT1G70782 | 198 | 6.57 | 25.71 | -688 | -0.63 | 0 | upstream open reading 28 (CPuORF) |
| AT3G11400 | 242 | 5.71 | 19.94 | -78 | -0.58 | 0 | eukaryotic translation initiation factor 2 sub.1 |
| AT5G43810 | 311 | 8.71348 | 49.17422 | -400 | -0.57 | 0 | Translation elongation factor EF1B γ-chain |
| AT2G40290 | 174 | 5.35016 | 17.60725 | -2405 | -0.51 | 0 | EIF2C (elongation initiation factor 2c) |
| AT1G04170 | 180 | 4.62 | 13.22 | -573 | -0.45 | 0 | eukaryotic translation initiation factor 2 sub- γ; |

**Supplemental Table 5**. LIMYB overexpression downregulates associated-genes photosystem II assembly, light reaction, and photosynthetic electron transport observed in the integrative RNA-seq/ChIP-seq analysis.

| Gene_id | width | Fold enrich. | log10 q-value | Distance to feature | log2 FC | padj | Description |
|---|---|---|---|---|---|---|---|
| AT1G15120 | 173 | 465 | 16.49 | -305 | -0.72 | 0 | Ubiquinol-cytochrome C reductase hing |
| AT1G16700 | 194 | 851 | 40.31 | -957 | -0.72 | 0 | Alpha-helical ferredoxin |
| AT1G75630 | 186 | 547 | 18.37 | -408 | -0.96 | 0 | vacuolar H+-pumping ATPase 16 kDa pro |
| AT4G04640 | 237 | 752 | 35.64 | -2103 | -0.76 | 0 | ATPase, F1 complex, gamma subunit pr |
| AT4G09650 | 400 | 1133 | 6.67 | -1043 | -1.33 | 0 | ATP synthase delta-subunit gene Encod |
| AT4G11150 | 189 | 62 | 23.18 | -1200 | -0.63 | 0 | vacuolar ATP synthase subunit E1 Encod |
| AT4G32260 | 331 | 1101 | 76.31 | -236 | -0.98 | 0 | ATPase, F0 complex, subunit B/B., bacte |
| AT5G40810 | 173 | 559 | 19.15 | -2033 | -0.07 | 0 | Cytochrome C1 family |
| AT5G40810 | 170 | 438 | 11.85 | -273 | -0.07 | 0 | Cytochrome C1 family |
| AT5G47890 | 234 | 58 | 21.81 | -1038 | -1.04 | 0 | NADH-ubiquinone oxidoreductase B8 s |
| AT5G47890 | 200 | 584 | 22.34 | -676 | -1.04 | 0 | NADH-ubiquinone oxidoreductase B8 s |
| AT5G66570 | 368 | 802 | 43.63 | -1109 | -0.07 | 0 | PS II oxygen-evolving complex 1 Encodes a p |
| AT1G06680 | 205 | 822 | 38.21 | -208 | -1.11 | 0 | photosystem II subunit P-1 |
| AT4G05180 | 300 | 681 | 28.34 | -878 | -0.59 | 0 | photosystem II subunit Q-2 |
| AT1G79040 | 170 | 486 | 14.64 | -586 | -1.34 | 0 | photosystem II subunit R |
| AT1G44575 | 229 | 651 | 25.96 | -1501 | -1.08 | 0 | Chlorophyll A-B binding family protein |
| AT2G30570 | 170 | 486 | 14.64 | -88 | -1.34 | 0 | photosystem II reaction center W Encodes P |
| AT1G67740 | 257 | 1155 | 66.25 | -1158 | -0.67 | 0 | photosystem II BY |
| AT1G52230 | 264 | 1153 | 68.15 | -2151 | -0.97 | 0 | photosystem I subunit H2 Phosphorylation o |
| AT1G10960 | 283 | 995 | 52.27 | -398 | -1.79 | 0 | ferredoxin 1 |
| AT1G20020 | 243 | 588 | 21.69 | -1150 | -1.16 | 0 | ferredoxin-NADP(+)-oxidoreductase 2 |
| AT4G04640 | 237 | 752 | 35.64 | -2103 | -0.76 | 0 | ATPase, F1 complex, gamma subunit protein |
| AT4G09650 | 400 | 1133 | 6.67 | -1043 | -1.33 | 0 | ATP synthase delta-subunit gene Encodes th |
| AT4G32260 | 331 | 1101 | 76.31 | -236 | -0.98 | 0 | ATPase, F0 complex, subunit B/B., bacterial/ |
| AT1G29910 | 372 | 1072 | 104.41 | -889 | -0.92 | 0 | chlorophyll A/B binding protein 3 membe |
| AT1G45474 | 206 | 62 | 23.18 | -964 | -0.13 | 0 | photosystem I light harvesting complex ge |
| AT3G27690 | 473 | 737 | 31.72 | -482 | -0.93 | 0 | photosystem II light harvesting complex g |
| AT3G47470 | 699 | 567 | 19.79 | -1654 | -1.03 | 0 | light-harvesting chlorophyll-protein comp |
| AT4G10340 | 336 | 854 | 40.96 | -291 | -0.88 | 0 | light harvesting complex of photosystem |
| AT4G10340 | 180 | 426 | 11.18 | -1234 | -0.88 | 0 | light harvesting complex of photosystem |
| AT5G54270 | 462 | 571 | 2.11 | -672 | -0.85 | 0 | light-harvesting chlorophyll B-binding prote |
| AT5G54270 | 217 | 492 | 17.51 | -2445 | -0.85 | 0 | light-harvesting chlorophyll B-binding p |

**Supplemental Table 6.** List of primers used for cloning, PCR and RT-qPCR

| Primer | Gene | Sequence 5'–3' |
|---|---|---|
| qRT- At2G21580 RPS25B -Fwd | At2g21580 (RPS25B) | CCGATCGTTACTCCGTCGAA |
| qRT- At2G21580 RPS25B -Rvs | At2g21580 (RPS25B) | AGATTTGGCCGGCTTTGAT |
| qRT- At3G07110 RPL13A -Fwd | At3g07110 (RPL13A) | GGCTGATCCAGAGCTGAGTGAAA |
| qRT- At3G07110 RPL13A -Rvs | At3g07110 (RPL13A) | GTGGTGACGAGCATCAACCA |
| qRT-Ef1B -Fwd | At5g12110 (Ef1B) | GTGGTACGATTCTGTTGCT |
| qRT-Ef1B -Rvs | At5g12110 (Ef1B) | CAGTATGTGGATGAGCTTCAG |
| qRT-eIF2α -Fwd | At5g05470 (eIF2α) | CTGCTCCGAATCTAGAATGTC |
| qRT-eIF2α -Rvs | At5g05470 (eIF2α) | ATACGTAAGCTCCCATGTCA |
| RT-qPCR Luci. Fire -Fwd | LOC116160065 (Luc. Firefly) | GCGAAGGTTGTGGATCTGGA |
| RT-qPCR Luci. Fire - Rvs | LOC116160065 (Luc. Firefly) | CGTCTTCGTCCCAGTAAGCT |
| RT-qPCR Actina -Fwd | At1g13180 (Actina) | ATGTCGTGAGCCATCCTGTC |
| RT-qPCR Actina -Rvs | At1g13180 (Actina) | ACACCGGATTCGTGCGGCAT |
| qRT-FD1 -Fwd | At1g10960 (FD1) | CAATCTCTCTTCGGCCTC |
| qRT-FD1 -Rvs | At1g10960 (FD1) | GTCGAGGACGTAGACATCT |
| qRT-PSIIOEC -Fwd | At2g28605 (PHSIIOEC) | AGCTTATCAACGCAGAGAA |
| qRT-PSIIOEC -Rvs | At2g28605 (PHSIIOEC) | TTGTTGTCCTAATCCTGCT |
| qRT-Hsp70 -Fwd | At3g12580 (HSP70) | GGCTGAGGCAGATGAGTTC |
| qRT-Hsp70 -Rvs | At3g12580 (HSP70) | GTGTCGTCATCCATTCCTC |
| qRT-RAB18 -Fwd | At5g66400(RAB18) | ACGAGTACGGAAATCCGATG |
| qRT-RAB18 -Rvs | At5g66400(RAB18) | ACCACCACTTTCCTTGTGGA |
| LIMYB At5g05800B - Fwd | At5g05800 (LIMYB) | AAAAAGCAGGCTTCACAATGAGGCCAAA AGCAGT |
| LIMYB At5g05800B - Rvs | At5g05800 (LIMYB) | AGAAAGCTGGGTCCTATGTTGTAGTAGG AATAGG |
| pRL18-TGW-Fwd (1) | AT1G29970 (RPL18)- -1000 pb promoter | GGGGACAACTTTGTATAGAAAAGTTGTC GGAAGATGAGTATACGTAAATA |
| pRL18-TGW-Rvs (1) | AT1G29970 (RPL18)- -1000 pb promoter | GGGGACTGCTTTTTTGTACAAACTTGCAA TTGAGATCCGATTAACCTAA |
| pRL18-TGW-Fwd (2) | AT1G29970 (RPL18)- -2000 pb promoter | GGGGACAACTTTGTATAGAAAAGTTGTC GTGTACCAGTAAAATGTTGTTG |
| pRL18-TGW-Fwd (3) | AT1G29970 (RPL18)- -325 pb promoter | GGGGACAACTTTGTATAGAAAAGTTGTC GACCATAATCATATTCATTGAAGTG |
| pPSIIOEC-TGW-Fwd (1) | At2g28605 (PHSIIOEC)- -2000 pb promoter | GGGGACAACTTTGTATAGAAAAGTTGTCGT CCCCTCCAATTCTGACA |
| pPSIIOEC-TGW-Rvs (1) | At2g28605 (PHSIIOEC)- -2000 pb promoter | GGGGACTGCTTTTTTGTACAAACTTGCTGC TTCTTTCTCCTTATCAAC |
| pFD1 TGW-Fwd (1) | At1g10960 (FD1) -2000 pb promoter | GGGGACAACTTTGTATAGAAAAGTTGTC GGACCCCTCCAGTGTTAC |
| pFD1 TGW-Rvs (1) | At1g10960 (FD1) -2000 pb promoter | GGGGACTGCTTTTTTGTACAAACTTGCTG TTGAAAGATGATTCTACAGA |
